# Axon initial segment as an axonal lysosome trafficking control gate

**DOI:** 10.64898/2026.11.20.768018

**Authors:** Yubing Li, Keyu Chen, Qiufeng Teng, Hongze Mao, Yaojun He, Fang Liu, Zhijie Lin, Rui Yang, Mingjie Zhang

**Affiliations:** Greater Bay Biomedical Innocenter, Shenzhen Bay Laboratory, Shenzhen 518132, China; Zhejiang Key Laboratory of Organ Development and Regeneration, College of Life and Environmental Sciences, Hangzhou Normal University, Hangzhou 311121, China; Institute of Neuroscience, Translational Medicine Institute, Health Science Center, School of Basic Medical Sciences, Xi’an Jiaotong University, Xi’an, 710061, China; School of Life Sciences, Southern University of Science and Technology, Shenzhen 518055, China

**Keywords:** Neuronal lysosomes, lysosomal quality-control, lysosomal axonal transport, axon initial segment (AIS), Ankyrin-G (*Ank3*), MCF2L

## Abstract

Spatial subcellular distributions of lysosomes are intricately coupled with their functional heterogeneities, yet how neurons preferentially allocate distinct subtypes of lysosomes between the soma and axons remains largely unknown. Here, we discover that the axon initial segment (AIS) may function as a lysosomal quality-control checkpoint by actively sorting lysosomes based on their functional states. Central to this checkpoint is MCF2L, a specific lysosome-associated protein enriched in AIS, that anchors clustered lysosomes at AIS via high-affinity binding to giant ankyrin-G (gAnkG, encoded by *Ank3*). Selective ablation of the MCF2L-gAnkG interaction accelerates disease progression in a mouse Parkinson’s disease model. By clustering lysosomes, MCF2L restricts kinesin-mediated transport and regulates lysosomal positioning. In neurons, MCF2L-mediated lysosome clustering may act as a lysosomal quality-control checkpoint, by which less acidic (e.g., impaired) lysosomes are retained in AIS and functionally intact lysosomes are released for transport. Thus, AIS may impart neurons with a unique mechanism for axonal lysosome compartmentalization and quality control.

## Introduction

Axons, comprising over 90% of neuronal volume and surface area, are the first neuronal regions compromised in degenerative diseases ^1^. Axons form the body’s slenderest structures ^2,3^, which at the same time are exposed to numerous cellular challenges including vulnerability to occlusions, proteotoxicities, metabolic dysfunctions, mechanical stresses, etc. Paradoxically, established early during development as permanent and irreplaceable entities, axons usually can maintain functional integrity over decades or even the entire lifetime of an animal ^4^. This remarkable resilience to cellular stresses implies that neurons must possess specialized mechanisms to safeguard the homeostasis of biomolecules and cellular organelles within the crowded axonal compartments ^4–9^. However, our understanding of the axonal functional homeostasis is still limited.

Lysosomes, which function not only as terminal degradative organelles but also as multifunctional signaling hubs, are crucial for maintaining neuronal homeostasis through orchestrating metabolic regulation, membrane repair, calcium signaling, and synaptic plasticity ^10–15^. Lysosomal dysregulation actively participates axonal impairments and constitutes a pivotal step in the pathogenesis of neurodegenerative diseases ^16–24^. Extensive studies in the past chiefly in non-neuronal cells revealed that subcellular distribution of lysosomes is tightly coupled with their subpopulation identities and functional heterogeneities ^25–27^. Though much less studied, neurons likely operate with a similar paradigm ^19,28^. Asymmetric distribution is a prominent feature of neuronal lysosomes: they predominantly reside in the soma and proximal dendrites, whereas only a minor fraction is present in axons ^28–32^. This distribution requires stringent regulation, as abundance of local active lysosomes influences the homeostasis of surrounding compartments ^28,33^. For example, excessive accumulation of immature or dysfunctional lysosomes may lead to axonal congestion or interfere with normal lysosomal function ^34–36^. However, mechanisms that govern the uneven distribution of distinct lysosomal subpopulations between the soma and axon remain poorly defined ^37,38^.

The polarized neuronal information flow requires strict compositional heterogeneity between axonal and somatodendritic compartments—a hallmark of neuronal polarity ensuring selective distributions of proteins, lipids and vesicular organelles ^39,40^. The axon initial segment (AIS), a specialized subcellular domain separating the soma and axon, is critical for maintaining neuronal polarity and axonal homeostasis ^41–44^. Defects in AIS lead to structural and functional impairments of axons and have been implicated in many neurodegenerative pathologies, including Alzheimer’s disease (AD) and amyotrophic lateral sclerosis (ALS) ^45–52^, but the underlying mechanisms are not fully understood. AIS maintains axon-dendrite polarity by restricting lateral diffusion of membrane proteins and selectively gating vesicular trafficking ^53–63^.

As the gateway for lysosomal shuttling between the axon and soma, the crosstalk between AIS and lysosomes remains unclear. Increasing evidence supports a potential role of AIS in the axonal transport of lysosomal vesicles. For instance, in motor neurons, lysosomes exhibit dynamic movement between the soma and AIS, with only a limited fraction entering the distal axon ^64^. At the pre-axonal region, which is closely associated with AIS, endoplasmic reticulum tubules facilitate lysosome fission, thereby supporting axonal transport and maintaining proper lysosome availability in axons ^32^. In addition, loss of TMEM106B, a lysosomal membrane protein linked to frontotemporal lobar degeneration, causes localized AIS swelling and abnormal accumulation of immature lysosomal vacuoles within the swollen structure ^65^, suggesting that AIS may participate in lysosomal quality control during lysosomal biogenesis and transport. However, the molecular mechanisms governing the trafficking of lysosomes between the soma and axons remain poorly understood.

In this study, we identify MCF2L (aka DBS, OST, or ARHGEF14) as a lysosomal membrane associated protein. In addition, MCF2L is also a critical molecular tether linking lysosomes to AIS via high-affinity binding to the AIS core scaffold protein giant ankyrin-G (gAnkG). Selective disruption of the gAnkG-MCF2L interaction via a structure-guided single-point mutation without affecting the overall structural integrity of AIS markedly accelerates pathology in a mouse Parkinson’s disease model. Mechanistically, lysosomes exhibit a distinct propensity to form clusters at AIS compared to other axonal compartments. This spatial lysosome clustering is driven by oligomerization of MCF2L following its gAnkG-binding-dependent enrichment at AIS. Once clustered at AIS, lysosomes lose their capacity for fast axonal transport. Importantly, less acidic lysosomes are preferentially retained in MCF2L-associated clustered at AIS, suggesting that AIS may function as a lysosome quality-control “checkpoint”.

## Results

### Searching for molecules bridging the axon initial segment and lysosomes identifies MCF2L as a novel lysosomal protein

In some neurons, lysosomes (labeled by endogenous Lamp1, LAMTOR4, or Magic Red) can be observed to be enriched at the AIS relative to other axonal regions (see Fig. S1 by Farías et al. **^66^**), implying that the AIS compartment may interact with lysosomes to regulate their distribution or transport. To evaluate this speculation, we osmotically stressed lysosomes/late endosomes (Ly/LE) in cultured hippocampal neurons using 100 mM sucrose, which induced swollen Ly/LE (labeled by Lamp1) **^67^**. Like native neurons, these swollen vesicles accumulated primarily in the soma, with a small portion distributed in the neurites. Interestingly, a substantial proportion of neurons also exhibited swollen Ly/LE trapped at AIS (Fig. 1A, 1B). These observations imply that AIS may be able to recognize functionally impaired lysosomes and restrict their passage to the distal axon or soma. Ankyrin-G (AnkG), specifically the 270 kDa or 480 kDa giant isoform (gAnkG), are major scaffold proteins of AIS. These giant AnkG isoforms are enriched exclusively at AIS and nodes of Ranvier, and the 480 kDa isoform is absolutely essential for AIS assembly and function **^68^**. Depletion of all AnkG isoforms causes complete disassembly of the AIS **^69,70^**, precluding observation of Ly/LE trapping at the AIS. We therefore replaced endogenous AnkG in neurons derived from *Ank3* exon 22/23-floxed mice with 480- or 270-kDa AnkG-GFP by co-transfection with Cre recombinase ⁶⁸ to determine whether gAnkG contributes to AIS-associated Ly/LE trapping. Expression of 480 kDa AnkG-GFP fully restored the sucrose-induced Ly/LE trapping at AIS, whereas 270 kDa AnkG-GFP could not (Fig. 1B). These results suggest that AIS is involved in regulating the axonal passage of lysosomes and that the long intrinsically disordered sequence unique to the 480 kDa AnkG isoform corresponding to the C-terminal portion of the neuronal specific domain (NSD-Cter, aa. 2329-4034; Fig. S1A), is indispensable for this function.

**Fig 1.**
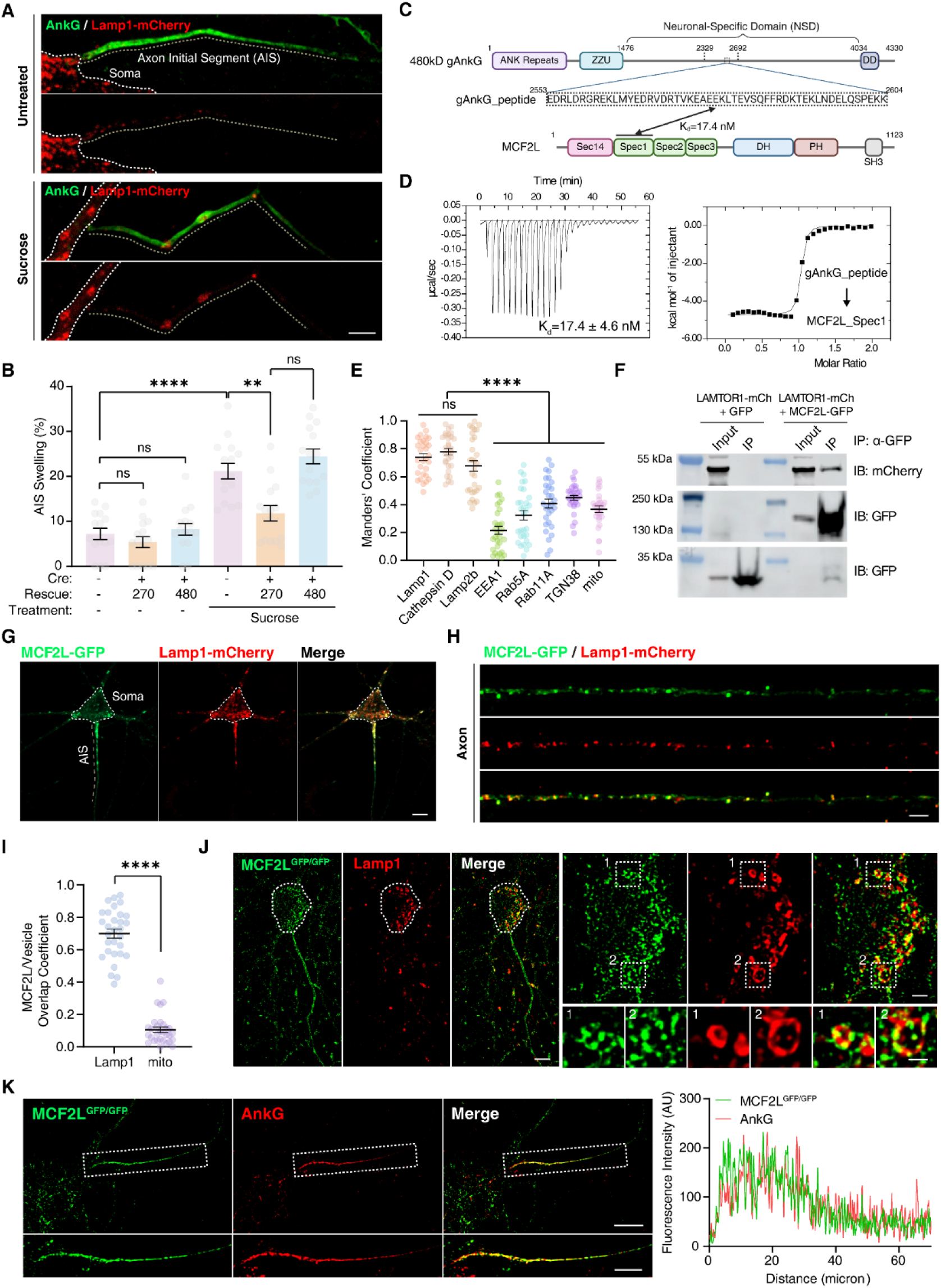
MCF2L is a lysosomal resident protein that is enriched at AIS via binding to gAnkG. **A.** Representative image showing sucrose-induced AIS swelling and lysosome clustering at AIS. Scale bar, 5 μm. **B.** Quantification of sucrose-induced AIS swelling in *Ank3*^flox/flox^ neurons expressing GFP as a control, or Cre together with rescue constructs encoding 270 kDa AnkG or 480 kDa gAnkG. Data are shown as mean ± SEM from three independent experiments; n=15 random fields of view. One-way ANOVA followed by Tukey’s post hoc test was used. ****p<0.0001, **p=0.0029; ns, not significant. **C.** Schematic diagrams of the domain organization of gAnkG and MCF2L. The minimal MCF2L-binding region of gAnkG and its amino acid sequence, referred to as the gAnkG_peptide (aa. 2553-2604), is indicated by the dashed box. **D.** ITC titration experiment quantitatively determines the binding affinity between gAnkG_peptide and WT MCF2L_Spec1. *Left*, raw titration curve; *right*, fitting curve. **E.** Manders’ colocalisation coefficients evaluating the overlap of MCF2L with different vesicle markers. One-way ANOVA followed by Tukey’s post hoc test was used. ****p<0.0001; ns, not significant. **F.** Co-immunoprecipitation of MCF2L-GFP or GFP (as a negative control) with LAMTOR1-mCherry in lysates of HEK293T cells co-transfected with both expression constructs. **G-H.** Exogenously expressed MCF2L-GFP displays a punctate pattern colocalizing with Lamp1 in neuronal soma and proximal dendrites (**G**) and in axons (**H**). Scale bars, 10 μm for panel **G** and 2 μm for panel **H**. **I.** Quantification of the colocalization of MCF2L with Lamp1 or the mitochondria marker mito in axons. Mean ± SEM, three independent experiments; n=30 axons. ****p<0.0001, two-tailed unpaired t test. **J.** Endogenous MCF2L localizes to lysosomes/late endosomes in neurons. *Left*: SIM images of a DIV7 cultured hippocampal neuron from MCF2L^GFP/GFP^ mouse immunostained for GFP and Lamp1. Scale bar, 5 µm. *Top right*: Magnified images of the somatic region of the selected neuron. Scale bar, 2 µm. *Bottom right*: Magnified images of selected lysosomes. Scale bar: 1 µm. **K.** Endogenous MCF2L localizes to AIS. SIM images of a DIV11 cultured hippocampal neuron from a homozygous MCF2L^GFP/GFP^ mouse immunostained for GFP and AnkG. Scale bar, 10 µm; magnified inset, 5 µm. Dashed box is selected for magnification (lower panel) and line-scan analysis (right panel).

We further hypothesized that the NSD-Cter of 480 kDa gAnkG interacts with lysosome-related proteins to mediate sucrose-induced Ly/LE trapping at AIS. Yeast two-hybrid screening using a recombinant fragment of this intrinsically disordered region (Fig. S1B) as a bait identified MCF2L (also known as DBS, OST or ARHGEF14), as the top candidate for interacting with gAnkG (aa. 2329-2691) (Fig. 1C). Both isothermal titration calorimetry (ITC) and size-exclusion chromatography coupled with multi-angle light scattering (SEC-MALS) confirmed that MCF2L strongly binds to gAnkG, with a dissociation constant (Kd) of ∼17 nM and a 1:1 binding stoichiometry (Fig. 1C, 1D, S1C). MCF2L is a multi-domain RhoGEF family member, containing an N-terminal Sec14 domain, three consecutive spectrin repeats (designated Spec1, Spec2, and Spec3), a DH domain (Dbl homology or RhoGEF domain), a PH domain (Pleckstrin homology domain), and an SH3 domain (Fig. 1C). Quantitative binding affinity-directed mapping by ITC revealed that the first spectrin repeat (MCF2L_Spec1) is necessary and sufficient for binding to gAnkG. Conversely, a minimal binding region of 52 amino acids (residues 2553-2604) within gAnkG NSD-Cter was sufficient for interacting with MCF2L (Fig. 1C, S1D). Interestingly, the corresponding binding sequences in gAnkG and MCF2L are highly conserved from chordates to human, coinciding with the evolutionary emergence of myelination and the dramatic increase in axon length of neurons (Fig. S2A, S2B), implying that the gAnkG-MCF2L interaction may have a conserved role in supporting the unique demands of long and myelinated axons.

MCF2L was originally identified from osteosarcoma and has been implicated in a diverse array of diseases, including osteoarthritis, osteoporosis, Alzheimer’s disease, and atherosclerosis **^71–75^**. However, its biological function and precise subcellular distribution remained poorly characterized. We first examined its colocalization with a panel of organellar markers in heterologous cells and primary cultured astrocytes, a CNS-derived cell type with an extended morphology that facilitates subcellular localization analysis. MCF2L exhibited strong colocalization with the Ly/LE marker Lamp1, Cathepsin-D, and Lamp2b, yielding mean Manders’ coefficients of approximately 0.7∼0.8 (Fig. 1E, S3A). In contrast, its colocalization with markers of early endosomes (EEA1 and Rab5A), recycling endosomes (Rab11A), the trans-Golgi network (TGN38), and mitochondria was markedly lower, with Manders’ coefficients in the range of 0.2∼0.4 (Fig. 1E, S3A).

MCF2L is preferentially expressed in the central nervous system compared with other tissues **^71,76,77^**. We then examined its localization in cultured hippocampal neurons. MCF2L-GFP displayed a punctate pattern along axons, the soma, and proximal dendrites, with pronounced colocalization with Lamp1 (Fig. 1G, 1H). Notably, a subset of MCF2L-positive structures formed larger clusters that were predominantly enriched in the proximal axon (Fig. 1G), indicating that MCF2L may mediate communication between AIS and lysosomes (see below for more details). By contrast, MCF2L showed poor colocalization with somatodendritic vesicles marked by transferrin receptor (TfR) or TGN38, and early endosomes marked by EEA1 or Rab5A (Fig. S3B), suggesting that MCF2L is also a lysosome-specific factor in neurons. SIM imaging further confirmed robust colocalization between MCF2L and Lamp1 in neurons (mean overlap coefficient: ∼0.7), in sharp contrast to the negative control using the mitochondrial marker COX8 (mito) (Fig. 1I and S3C). Live-cell imaging also revealed rapid cotransport of MCF2L-GFP with Lamp1-positive vesicles in the proximal axonal regions (Movie 1). Consistent with these findings, co-immunoprecipitation assays detected the lysosomal marker LAMTOR1-mCherry in MCF2L-GFP immunoprecipitates (Fig. 1F). Together, these results identify MCF2L as a lysosome-associated protein in neurons and non-neuronal cells.

To identify the lysosomal membrane-targeting region, we generated truncation mutants of MCF2L and assessed their colocalization with Lamp1 (Fig. S4A, S4B). The Sec14 and Spec1 domains together were necessary and sufficient for MCF2L to colocalize with lysosomes (Fig. S4B, S4C). Deletion of either domain resulted in diffused cytoplasmic distribution of the mutant MCF2L (Fig. S4B). The Sec14 and Spec1 domains are separated by only a very short (3 amino acids) and conserved linker (Fig. S2B), hinting that the two domains may function as a structurally coupled supramodule. Supporting this, AlphaFold3-predicted structure shows tight packing of Sec14 and Spec1 into a compact supramodule (Fig. S4D). Based on this predicted structure, we substituted Tyr85 in the Sec14 domain with Gln (Y85Q) or Trp195 in Spec1 with Arg (W195R). These point mutations were expected to destabilize or even disrupt the interaction between the Sec14 and Spec1 domains in the supramodule (Fig. S4E). As expected, MCF2L carrying either point mutation displayed a largely diffused cytoplasmic localization and failed to colocalize with Lamp1 (Fig. S4E, S4F). Together, these results demonstrate that the tightly coupled Sec14-Spectrin supramodule is responsible for MCF2L to associate with lysosomes.

To validate the subcellular localization of endogenous MCF2L, we generated a knock-in (KI) mouse strain expressing an EGFP tag fused to the carboxy terminus of MCF2L (MCF2L^GFP/GFP^, to be differentiated from the MCF2L-GFP overexpression construct used throughout the study) as reliable commercial or custom-made antibodies suitable for immunocytochemistry were unavailable/unsuccessful. SIM imaging of hippocampal neurons derived from this KI mouse strain revealed that endogenous MCF2L exhibited a punctate distribution in both the soma and neurites (Fig. 1J), with a subset of these puncta coalescing into amorphous aggregated clusters within a single proximal neurite—consistent with observations in neurons exogenously expressing MCF2L-GFP (Fig. 1J vs 1G). Endogenous MCF2L signals in this neurite demonstrated extensive colocalization with AnkG, indicating enrichment of MCF2L at AIS (Fig. 1K). In addition, closer examination of magnified MCF2L puncta images revealed that endogenous MCF2L formed small patches decorating the lysosomal surface (Fig. 1J, lower right panels). In summary, these results reveal that MCF2L associates with lysosomal membranes in neurons.

### gAnkG enriches MCF2L at AIS

As most currently known AIS proteins depend on gAnkG for their localization to AIS ^78^, we therefore asked whether this also applies to MCF2L. Considering that global knockout or knockdown of gAnkG inevitably disrupts AIS assembly and complicates data interpretation, we sought to identify point mutations in gAnkG or MCF2L that specifically disrupt their interaction while preserving the functions of each protein.

We crystallized MCF2L_Spec1 (amino acids 196-326) in complex with gAnkG_peptide (aa. 2553-2604). The crystal structure of the complex, determined at a resolution of 2.9 Å, revealed a 1:1 binding mode, consistent with the ITC/SEC-MALS results (Fig. 1D, 2A and S1C). MCF2L_Spec1 adopts a canonical three-helix spectrin-repeat fold, and the gAnkG_peptide forms a single α-helix binding to MCF2L_Spec1, completing a four-helix bundle structure (Fig. 2A). The binding interface of the complex is predominantly hydrophobic and is further stabilized by charge-charge interactions (Fig. 2A, 2B). Such interactions presumably contribute to the high affinity and specificity of the complex formation. Interface residues are highly conserved in both proteins across the evolution (Fig. 2C). Within this interface, MCF2L Phe307, together with Leu216, and Phe304, forms a hydrophobic pocket that snugly accommodates Phe2587 from gAnkG (Fig. 2A, 2B). Substituting MCF2L Phe307 with the hydrophilic residue glutamine (F307Q) abrogated the interaction (Fig. 2D, upper panel) without altering the overall folding of MCF2L_Spec1 (Fig. S5A, S5B). Reciprocally, replacing gAnkG Phe2587 with glutamine (F2587Q) also eliminated its binding to MCF2L_Spec1 (Fig. 2D, lower panel). Since Phe2587 is located in the long and disordered region of gAnkG, this mutation is unlikely to perturb the overall structure of gAnkG. Together, these structural and biochemical studies elucidate the binding model and provide specific single point mutants in gAnkG and MCF2L, respectively, for dissecting the neuronal functions of these two large, multi-domain proteins and their interaction using cultured neurons and mice, as described below.

**Fig 2.**
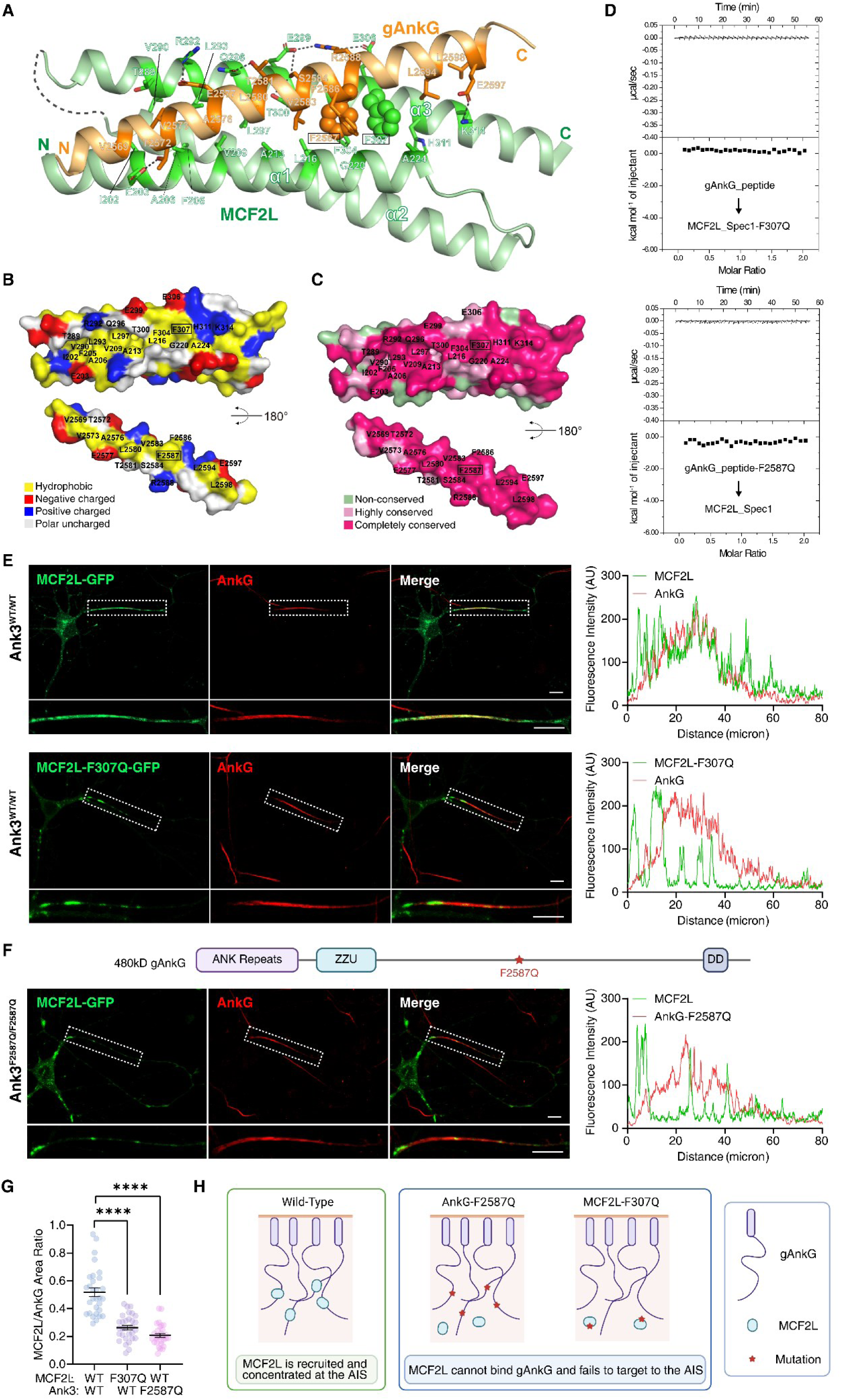
Structural basis of the gAnkG-MCF2L interaction and its role in MCF2L localization at AIS. **A.** Ribbon diagram of MCF2L_Spec1 in complex with gAnkG_peptide. Residues in the interaction interface are labeled. MCF2L F307 (green) and AnkG F2587 (orange) at the interaction core are shown as spheres. **B-C.** Open-book view of the MCF2L_Spec1/gAnkG_peptide complex shown in the surface charge distributions (**B**) and the amino acid residue conservation levels (**C**). **D.** ITC curves showing the binding of WT gAnkG_peptide to the MCF2L_Spec1-F307Q mutant (upper panel), and gAnkG_peptide-F2587Q to the WT MCF2L_Spec1 (lower panel). **E-F.** MCF2L localization at AIS depends on its binding to gAnkG. Confocal images of cultured hippocampal neurons expressing MCF2L-GFP or MCF2L-F307Q-GFP (**E**), or neurons from *Ank3*^F2587Q/F2587Q^ mice expressing MCF2L-GFP (**F**), immunostained for AnkG. Scale bars, 10 µm. Dashed boxes are selected for magnification and line-scan analysis. **G.** Quantification of overlaps (in areas) between MCF2L and AnkG at AIS. Mean ± SEM, three independent experiments; n=30 AISs. ****p<0.0001, two-tailed unpaired t test. **H.** Cartoon illustration showing that abolishing the interaction between gAnkG and MCF2L disrupts the enrichment of MCF2L at AIS.

To determine whether gAnkG-MCF2L binding determines MCF2L enrichment at the AIS, we first expressed the binding-deficient mutant MCF2L-F307Q-GFP, a mutant that displayed a normal lysosome binding property (Fig. S5C), in cultured neurons. Unlike wild-type (WT) MCF2L, the mutant exhibited markedly reduced AIS signals and diminished AnkG colocalization, retaining only occasional puncta at AIS (Fig. 2E, 2G). This indicates that AIS enrichment of MCF2L requires its direct binding to gAnkG. To further validate this *in vivo*, we generated an *Ank3*^F2587Q/F2587Q^ knock-in mouse bearing a homozygous F2587Q single-point mutation in gAnkG (Fig. S5D; see ***Methods*** for the designing details). This gAnkG F2587Q point mutation selectively disrupted its binding to MCF2L without altering the gAnkG-mediated AIS assembly. Consistent with this, the AIS structure in neurons from *Ank3*^F2587Q/F2587Q^ knock-in mice remained intact (i.e., the core AIS assembly function of gAnkG was unaltered; Fig. 2F). When MCF2L-GFP was expressed in *Ank3*^F2587Q/F2587Q^ neurons, it formed puncta in the soma and neurites but failed to enrich at AIS or to colocalize extensively with AnkG; only sparse MCF2L puncta were occasionally observed within the AIS region (Fig. 2F, 2G). Collectively, these results demonstrate that MCF2L localization to AIS and its extensive colocalization with AnkG strictly depend on the direct gAnkG-MCF2L interaction (Fig. 2H).

### Abolishing the MCF2L-gAnkG interaction accelerates neurodegenerative disease progression

Before further dissecting the roles and mechanisms of MCF2L in axonal lysosome trafficking, we first asked whether the gAnkG-MCF2L interaction is functionally important for neurons both in cultured neurons and mice. Although the gAnkG-MCF2L interaction is abolished, homozygous *Ank3*^F2587Q/F2587Q^ mice are viable and fertile, exhibiting no overt abnormalities in appearance, growth, or behavior compared with wild-type littermates (Fig. S6A-S6F). Cultured hippocampal neurons from mutant and wild-type control mice also showed no AIS morphological differences over days to weeks (Fig. S6G-S6L). These data indicate that this specific interaction is dispensable for AIS assembly, mouse development, and survival into adulthood under standard housing conditions.

We hypothesized that the phenotypic consequences of abolishing the gAnkG-MCF2L interaction might not be apparent under normal physiological conditions but could become evident under pathological stress. To test this, we used expression of the human Parkinson’s disease (PD)-associated α-synuclein mutant (hSNCA-A53T) as a cellular stressor. hSNCA-A53T promotes the formation of toxic α-synuclein fibrils (Lewy neurites), which are enriched in axons and ultimately contribute to axonal degeneration **^79,80^**. Pathogenic α-synuclein aggregates/fibrils can accumulate within neuronal lysosomes and are primarily cleared through lysosomal degradation **^81^**. We expressed the hSNCA-A53T using adeno-associated viruses (AAV). Notably, *Ank3*^F2587Q/F2587Q^ neurons expressing hSNCA-A53T exhibited larger aggregated α-synuclein puncta and greater average fluorescence intensity of these Lewy neurites than the wild-type controls (Fig. 3A-3C). These results indicate that selective disruption of the gAnkG-MCF2L interaction exacerbates axonal α-synuclein pathology, consistent with impaired clearance of pathogenic α-synuclein aggregates in axons.

**Fig 3.**
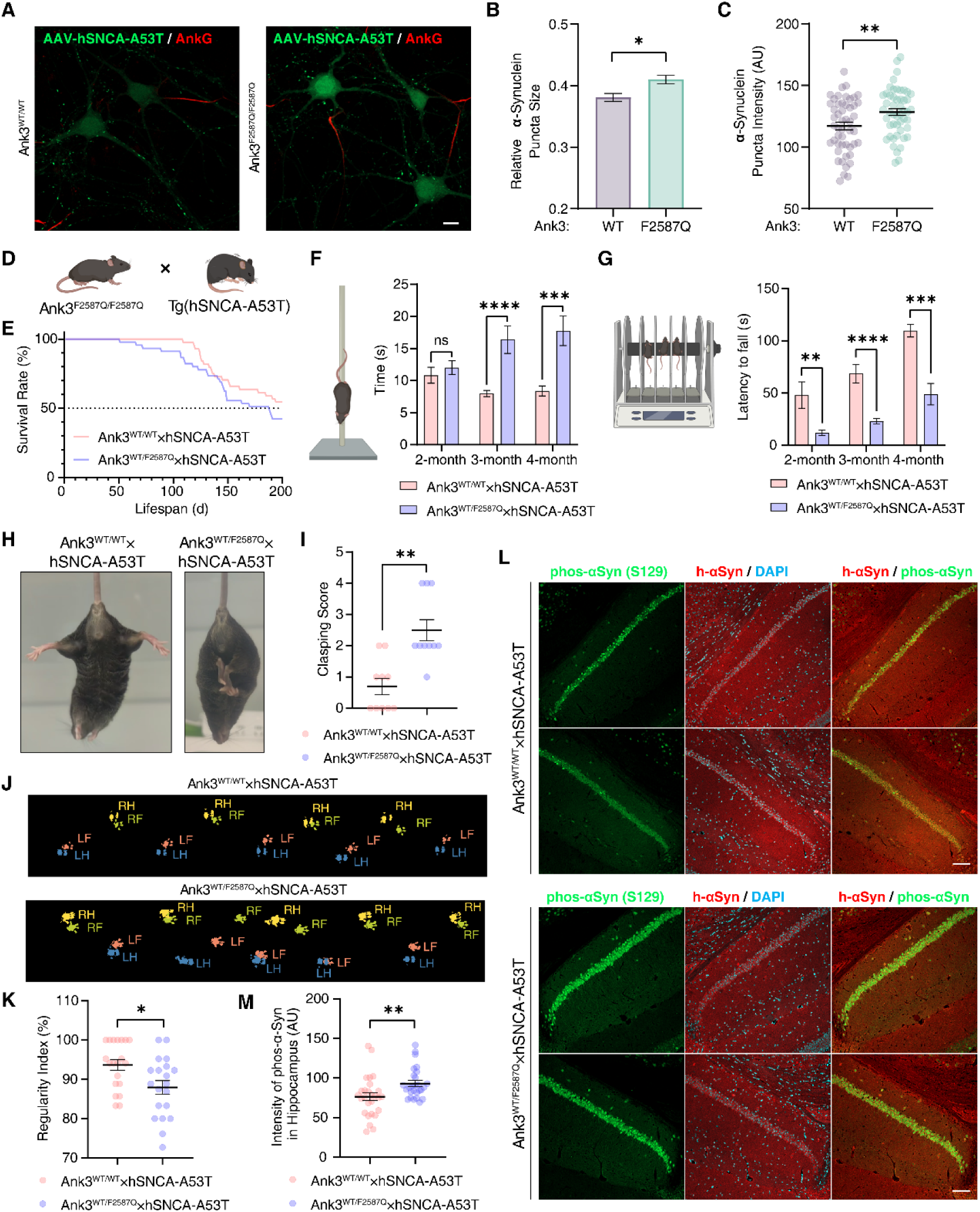
Disruption of the gAnkG-MCF2L interaction exacerbates α-synuclein accumulation and motor deficits in hSNCA-A53T mice. A-C. α-synuclein aggregates/fibrils accumulate more heavily in *Ank3*^F2587Q/F2587Q^ neurons. **A.** Images of neurons infected with AAV expressing α-synuclein mutant and immunostained for AnkG and α-synuclein. Scale bar, 10 μm. Quantification of α-synuclein puncta size (**B**) and intensity (**C**) in *Ank3*^WT/WT^ and *Ank3*^F2587Q/F2587Q^ neurons. Mean ± SEM, three independent experiments; n=4000 puncta (**B**) or 50 puncta (**C**). *p=0.0476, Mann-Whitney test; **p=0.0071, two-tailed unpaired t test. **D.** Breeding strategy used to generate *Ank3*^WT/WT^×hSNCA-A53T and *Ank3*^WT/F2587Q^×hSNCA-A53T mice. **E.** Kaplan-Meier survival curves for male *Ank3*^WT/WT^×hSNCA-A53T and *Ank3*^WT/F2587Q^×hSNCA-A53T mice; n=37 and n=35 mice, respectively. **F-G.** Motor performance of male *Ank3*^WT/WT^×hSNCA-A53T and *Ank3*^WT/F2587Q^×hSNCA-A53T mice in the pole test (**F**) and rotarod test at 10 rpm (**G**). Mean ± SEM, three independent experiments; **F**, n=12 mice; ***p=0.0009, two-tailed unpaired Student’s t test; ****p<0.0001, Mann-Whitney test. **G**, n=9, 19, 10 mice at 2, 3, and 4 months, respectively; **p=0.0036, ***p=0.0001, and ****p<0.0001, Mann-Whitney test. **H-I.** Representative limb-clasping phenotypes in 8-month-old mice (**H**) and quantification of clasping scores (**I**). Mean ± SEM, three independent experiments; n=10 mice. **p=0.0036, and **p=0.0013, Mann-Whitney test. **J-K.** CatWalk gait analysis showing representative paw-print patterns (**J**) and quantification of the regularity index (**K**) in *Ank3*^WT/WT^×hSNCA-A53T and *Ank3*^WT/F2587Q^×hSNCA-A53T mice. Mean ± SEM, three independent experiments; n=20 trials. *p=0.015 and **p=0.0013, Mann-Whitney test. **L-M.** α-synuclein staining in the hippocampal CA1 region of 3-month-old mice. **L.** Sections were immunostained for human α-synuclein (red), phospho-α-synuclein Ser129 (green) and DAPI (blue). Scale bar, 100 μm. **M.** Quantification of phospho-α-synuclein intensity. Mean ± SEM, three independent experiments; n=20. *p=0.0263, Mann-Whitney test.

To extend these findings *in vivo*, we crossed human α-synuclein A53T-mutant (hSNCA-A53T) transgenic mice, a well-established PD model, with homozygous *Ank3*^F2587Q/F2587Q^ mice to generate F1 *Ank3*^WT/F2587Q^ × hSNCA-A53T progeny (Fig. 3D, also see ***Methods*** and S7A). Unexpectedly, attempts to obtain homozygous *Ank3*^F2587Q/F2587Q^ mice carrying the hSNCA-A53T allele yielded very few animals, even after extended breeding trials, deviating markedly from Mendelian inheritance, and rendering functional characterization impractical (Fig. S7B). This result suggests that homozygous disruption of the gAnkG-MCF2L interaction is poorly tolerated in the context of α-synuclein disease burden. We therefore focused subsequent characterizations on viable *Ank3*^WT/F2587Q^ × hSNCA-A53T mice, using male mice for the experiments. Compared with *Ank3*^WT/WT^ × hSNCA-A53T littermates, the mutant (*Ank3*^WT/F2587Q^ × hSNCA-A53T) mice exhibited comparable body weight gain from 4 to 12 weeks of age (Fig. S7C), indicating grossly normal early developmental growth. However, Kaplan-Meier survival analysis revealed a significantly shortened lifespan in the mutant strain (Fig. 3E). These mice also developed more severe motor and sensory deficits at comparable ages (Fig. 3F-3K and Fig. S7D-S7G). In the pole test, no difference was observed at 2 months of age, but by 3 and 4 months, mutant mice required approximately twice as long to descend as control littermates (Fig. 3F), suggesting impaired movement initiation, motor flexibility or coordination. Similarly, in the rotarod test—a commonly used assay for neuronal dysfunction-induced motor deficits—*Ank3*^WT/WT^ × hSNCA-A53T mice remained on the rotating rod nearly twice as long as *Ank3*^WT/F2587Q^ × hSNCA-A53T mice (Fig. 3G), indicating profound impairments in motor learning, coordination, balance, and endurance. Hindlimb clasping, a neurological hallmark of disease progression in neurodegenerative models including Parkinson’s disease, was scored significantly higher in *Ank3*^WT/F2587Q^ × hSNCA-A53T mice (Fig. 3H, 3I). Gait analysis further supported accelerated locomotor dysfunction in mutant mice. Representative footprint patterns revealed disrupted gait organization in *Ank3*^WT/F2587Q^ × hSNCA-A53T mice (Fig. 3J). Quantitatively, the regularity index, which measures inter-paw coordination, was markedly reduced in the mutant group (Fig. 3K), reflecting impaired interlimb coordination and gait symmetry that point to underlying neuronal deficits. Consistent with this, additional gait parameters, including swing phase duration, stride length, single stance time, and body speed variation, were also significantly altered in the mutant mice (Fig. S7D-S7G). Together, these behavioral analyses demonstrate that even partial disruption of the gAnkG-MCF2L interaction accelerates neurological declines in the hSNCA-A53T mouse disease model.

Consistent with the behavioral phenotypes, immunohistochemical analysis revealed elevated levels of phosphorylated α-synuclein, a pathological hallmark of PD progression, in the brains of *Ank3*^WT/F2587Q^ × hSNCA-A53T mice compared with their *Ank3*^WT/WT^ × hSNCA-A53T littermates (Fig. 3L, 3M). Notably, total hSNCA-A53T protein levels were comparable between the two genotypes (Fig. S7H), indicating that a similar mutant human α-synuclein load produced greater accumulation of phosphorylated α-synuclein (the sum of human and mouse species) in *Ank3*^WT/F2587Q^ mice. Collectively, these data demonstrate that heterozygous disruption of the MCF2L-gAnkG interaction is associated with impaired clearance of toxic α-synuclein aggregates and accelerated disease progression, consistent with a role for lysosomal functions in axons **^81–83^**.

### MCF2L induces lysosomal clustering via dynamic oligomerization

To elucidate how the MCF2L-gAnkG interaction modulates lysosomal function, we first investigated whether MCF2L affects lysosomal behaviors. The coalescence of MCF2L-GFP clusters with Lamp1-mCherry positive clusters in neurons (Fig. 1G) suggested that MCF2L may have an intrinsic capacity to cluster Ly/LEs. To substantiate this, we expressed MCF2L-GFP in HeLa cells and observed a striking aggregation of Lamp1-labeled vesicles, in stark contrast to control cells transfected with GFP alone (Fig. S8A, S8B). To further establish this organelle-clustering capability of MCF2L in neurons, we conducted a gain-of-function assay utilizing a rapamycin-inducible heterodimerization system. TGN38, a trans-Golgi network (TGN)-specific integral membrane protein that exhibits no intrinsic colocalization with MCF2L or lysosomes, was fused to mCherry-FKBP tag and co-expressed with MCF2L-GFP-FRB in neurons (Fig. S8C). Without rapamycin, TGN38 remained scattered throughout the soma. Upon rapamycin-induced FKBP-FRB dimerization, TGN38-mCherry-FKBP labeled vesicles rapidly coalesced into clusters within 10 min, extensively colocalizing with MCF2L and were frequently positioned in close apposition to lysosomes (Fig. S8D, S8E). These results directly demonstrate the robust organelle clustering capacity intrinsic to MCF2L and also underscore MCF2L’s capability of tightly associating with lysosomes. This TGN-lysosome clustering assay also indicates that MCF2L associated with organelles can dimerize in-trans to cluster these organelles.

Notably, we observed a clear positive correlation between MCF2L expression levels and the formation of large lysosomal clusters, suggesting a concentration-dependent clustering mechanism (Fig. S8F, S8G). To understand the molecular basis of this phenomenon, we characterized the oligomerization dynamics of MCF2L *in vitro*. Recombinant MCF2L_Spec2-3 (encompassing the Spec2 and Spec3 domains) exhibited concentration-dependent dimerization, indicating a dynamic monomer-dimer equilibrium (Fig. 4A, 4B). MCF2L_Spec1-3 displayed similar features, but MCF2L_Spec1-2 existed as a stable monomer, indicating that the Spec3 domain mediates dimerization (Fig. S8H). Consistent with this result, AlphaFold3-based structural prediction showed that MCF2L_Spec1-3 adopts a parallel, symmetric, V-shaped dimer mediated by the Spec3 domain (Fig. 4C). In the predicted Spec3 dimer interface, Arg518 and Ser521 form a string of hydrogen bonds stabilizing association between the two spectrin repeats (Fig. 4C). To validate this structural model, we engineered an R518A/S521E double mutant (i.e., simultaneously substituting Arg518 with Ala and Ser521 with Glu) of MCF2L_Spec1-3 designed to disrupt the predicted Spec3 dimer interface. SEC-MALS analysis confirmed that the Spec1-3 mutant remained monomeric even at concentrations as high as 150 μM (Fig. 4D).

**Fig 4.**
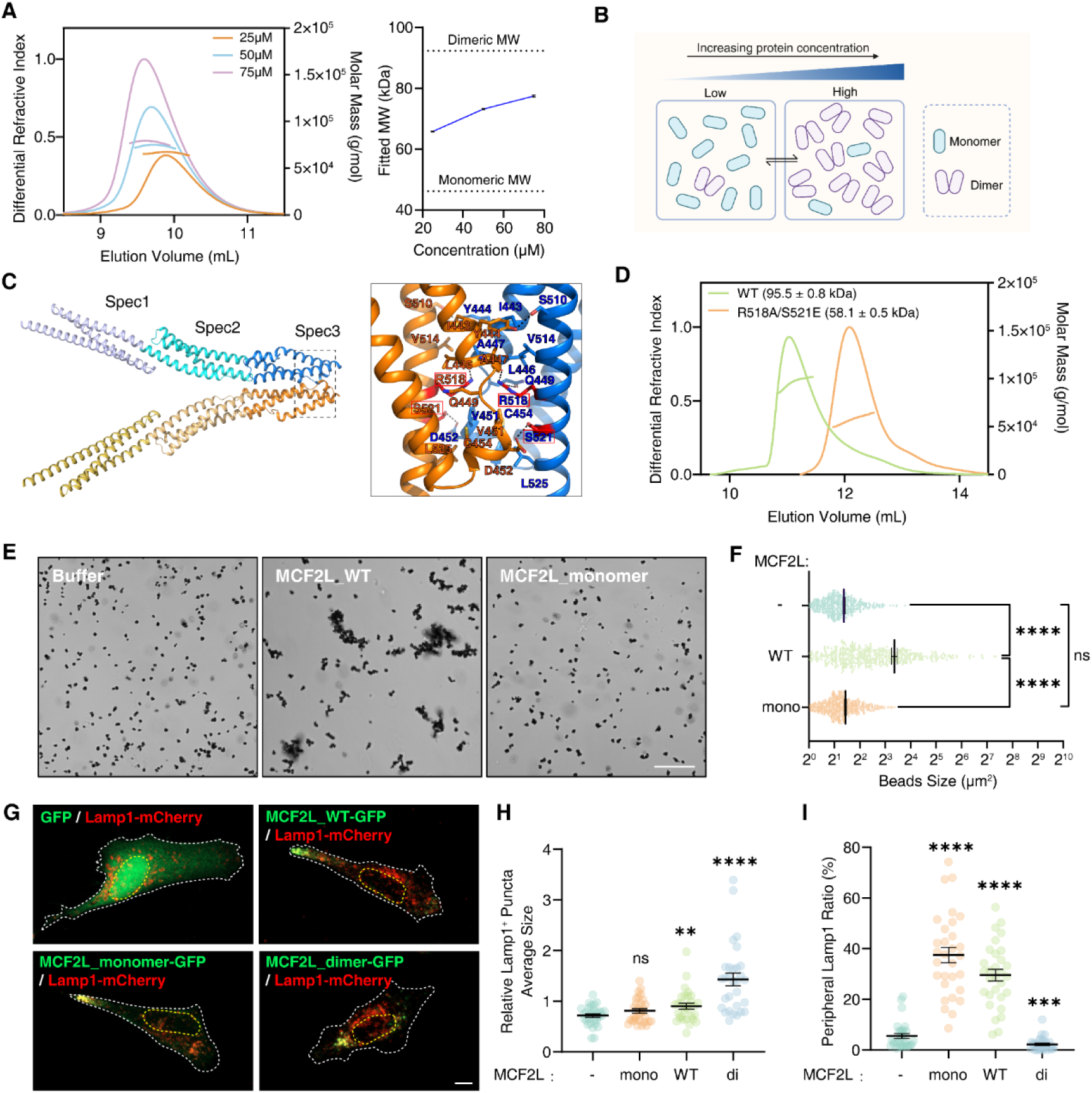
MCF2L induces lysosomal clustering via dynamic oligomerization. A-D. MCF2L dimerization is mediated by its Spec3 domain. **A.** SEC-MALS analysis of MCF2L_Spec2-3 at different concentrations and quantification of concentration-dependent apparent molecular-weight changes; mean ± SEM. **B**. Schematic representation of the dynamic monomer-dimer equilibrium of MCF2L. **C.** AlphaFold3-predicted structure of the MCF2L_Spec1-Spec3 dimer, with a magnified view showing the dimerization interface. Critical residues R518 and S521 are boxed. **D.** SEC-MALS analysis and molecular-weight comparison of WT and R518A/S521E MCF2L_Spec1-3 proteins at the loading protein concentration of 150 μM; mean ± SEM. **E-F.** *In vitro* bead clustering by MCF2L. **E.** Representative images of streptavidin-coated nano-beads incubated with StrepII-tagged WT MCF2L or the monomeric mutant. Scale bar, 20 μm. **F.** Quantification of bead clustering after incubation with buffer or the indicated MCF2L proteins. Mean ± SEM, three independent experiments; n=500 beads. ****p<0.0001; ns, not significant, Mann-Whitney test. **G-I.** Oligomerization states of MCF2L determine the cluster sizes and the subcellular positions of the protein itself and lysosomes. Representative images (**G,** scale bar: 10 μm; images of individual channels are shown in Figure S9A), quantifications of Lamp1-positive puncta average size (**H**), and quantifications of Lamp1-positive puncta distributions (**I**) in cells transfecting WT, monomeric, dimeric MCF2L or GFP. For **H**, n=30 cells; mean ± SEM. ****p<0.0001, **p=0.0073; ns, not significant, Mann-Whitney test. For **I**, n=30 cells; mean ± SEM. ****p<0.0001, ***p=0.0004, Mann-Whitney test.

We next tested whether MCF2L dimerization is sufficient to promote clustering in a reconstituted system. Microbeads with approximately 1 μm in diameter were functionalized with StrepII-tagged MCF2L proteins (Spec1-3) and used as lysosome mimics. WT MCF2L robustly aggregated beads into large agglomerates, whereas the monomeric R518A/S521E mutant lacked this activity, confirming that MCF2L dimerization can promote bead clustering (Fig. 4E, 4F). The assay further confirms that at least a certain proportion of MCF2L can dimerize in-trans (i.e., two MCF2L molecules from different beads) to promote bead clustering. To further assess the functional importance of dimerization, we also engineered a constitutively dimeric MCF2L_Spec1-3 variant (denoted as “locked-dimer” or “MCF2L_dimer”) by fusing the well-characterized GCN4 leucine zipper dimer **^84^** to the monomeric mutant (R518A/S521E). SEC-MALS confirmed that this chimeric protein existed as stable dimers across a broad concentration range (Fig. S8I). These engineered variants provided tools to compare the effects of reversible, abolished, or enforced MCF2L dimerization on lysosome clustering and distribution. Expression of monomeric MCF2L did not significantly alter the average size of lysosomal puncta compared with control cells. In contrast, WT MCF2L increased the average puncta size, indicative of enhanced lysosomal clustering, and the locked-dimer mutant induced much more pronounced lysosomal aggregation (Fig. 4G, 4H and S9A). In summary, these findings indicate that MCF2L potently clusters lysosomes through its dimerization capacity.

### Clustering of lysosomes changes their subcellular distribution

Interestingly, WT and monomeric MCF2L puncta frequently accumulated at cell tips and corners in nonneuronal cells, where they colocalized with Lamp1 (Fig. 4G, 4I and S9A). In contrast, the locked-dimer mutant showed no such enrichment at cell tips or corners, with most puncta instead localizing to the perinuclear region (Fig. 4G, 4I and S9A). This observation suggests that MCF2L is capable of modulating lysosome positioning in cells. To further substantiate this finding, we employed the FKBP-FRB system to artificially tether EEA1—a marker of early endosomes that normally do not accumulate at cell peripheries—to MCF2L. Upon rapamycin treatment, EEA1-mCherry-FKBP and MCF2L-GFP-FRB became colocalized, and a substantial fraction of MCF2L-tethered EEA1 was directed to the cell periphery, in stark contrast to the controls (Fig. S9B, S9C).

Domain-mapping experiments showed that deletion of the SH3 domain, the DH-PH domains, or even the entire C-terminal half of MCF2L, leaving only the Sec14 and Spec1-3 region, did not alter its cell peripheral enrichment (Fig. 5A, 5B). Consistent with the observations in Fig. 4G, 4I, the monomeric mutant of MCF2L accumulated predominantly at the cell periphery, whereas the locked-dimer mutant was almost exclusively concentrated in the perinuclear region (Fig. 5A, 5B). These findings indicate that the distribution of MCF2L at the cell periphery is governed by its oligomeric state.

**Fig 5.**
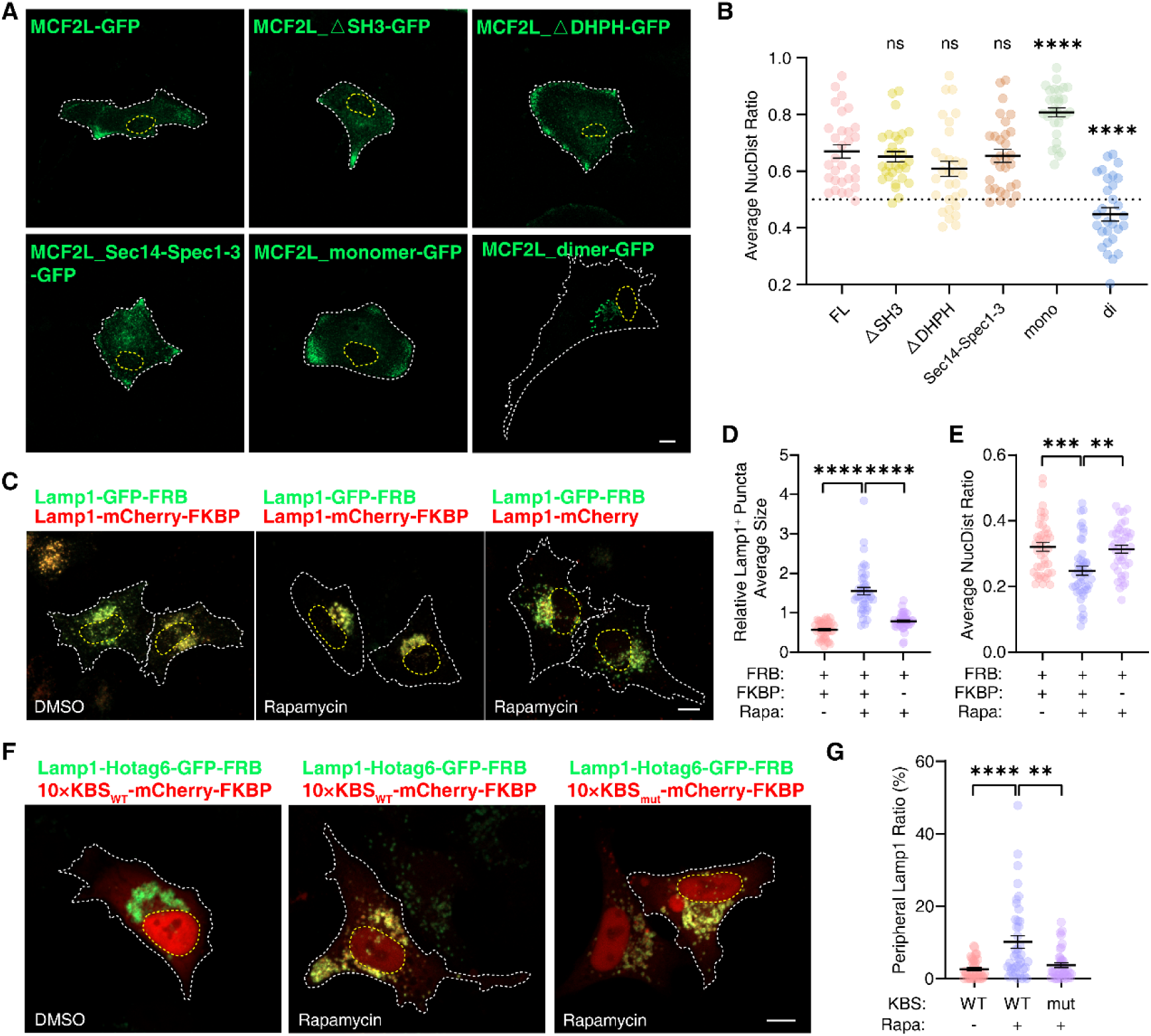
MCF2L oligomerization promotes lysosome clustering and positioning in perinuclear regions A-B. Mapping of the domain in MCF2L required for its localization to the cell cortex. **A.** Representative images showing distributions of different MCF2L truncation or deletion mutants in heterologous cells. **B.** Quantification of the average NucDist value for each mutant (see *Methods*). Mean ± SEM, three independent experiments; n=30 cells. ****p<0.0001; ns, not significant, two-tailed unpaired t test. **C-E.** Rapamycin-induced FRB-FKBP heterodimerization forces dimerization of Lamp1, leading to clustering of lysosomes to the perinuclear regions. **C**, representative images of cells expressing the indicated proteins with or without rapamycin. Scale bar, 10 μm.; **D**, Quantification of the average size of Lamp1-positive puncta. n =40 cells; mean ± SEM. One-way ANOVA followed by Tukey’s post hoc test. ****p < 0.0001. **E.** Quantification of the average NucDist value. n =40 cells; mean ± SEM. One-way ANOVA followed by Tukey’s post hoc test. ***p =0.0003, **p=0.0011. **F-G.** Clustered lysosomes can still be driven to the cell cortex by sufficient kinesin motors. **F**, representative images of cells expressing the indicated proteins with or without rapamycin. “10×KBS” indicates 10 copies of short kinesin-binding sequences chained in tandem. **G**, quantifications of cell peripheral Lamp1 fraction. n =40 cells; mean ± SEM. One-way ANOVA followed by Tukey’s post hoc test was used. ****p<0.0001, **p=0.0085.

We hypothesized that oligomerization of lysosomal membrane/membrane-associated proteins is sufficient to drive lysosomal clustering, impede lysosomal anterograde transport and consequently cause perinuclear retention. To test this, Lamp1 was fused to either FRB-GFP or FKBP-mCherry, and both constructs were co-expressed in cells. Rapamycin-induced crosslinking of Lamp1, which mimics protein oligomerization on the lysosomal membrane surface, not only induced the formation of large lysosomal clusters but also caused lysosomes to slowly retract from the cell periphery to the perinuclear region (Fig. 5C-5E and Movie 2). This finding was further corroborated by a parallel gain-of-function assay, in which MCF2L_Spec2-3 (in WT, monomer, or locked-dimer configurations) was fused to Arl8b, a lysosomal membrane-associated protein ^85^, which served as a lysosome-anchoring module. The monomeric Arl8b-Spec2-3 fusion construct had no significant effect on lysosomal distribution when compared with Arl8b, whereas the WT and locked-dimer constructs induced progressively stronger perinuclear accumulation (Fig. S9D, S9E).

One possible explanation for the impaired peripheral distribution of clustered lysosomes is that the clusters are too large to be efficiently transported by the limited number of kinesin molecules available on each lysosome. To test this, we first induced Lamp1 oligomerization by fusing them to HoTag6-GFP-FRB, which caused lysosome clustering and accumulation at the perinuclear region. Linking high concentrations of kinesins to these clusters by rapamycin-mediated tethering of a ten-copy kinesin-binding sequence (KBS) fused to mCherry-FKBP (10 **×**KBS-mCherry-FKBP) to Lamp1 successfully drove some of the large clusters to the cell periphery (Fig. 5F, 5G). Whereas the negative control containing a mutant version of KBS that cannot interact with kinesins (10×KBS_mut_-mCherry-FKBP) did not show such effects (Fig. 5F, 5G). These data indicate that lysosome clustering impedes kinesin-mediated transport of lysosomes to the cell periphery. Taken together, our results demonstrate that oligomerization of lysosomal membrane associated proteins (e.g., MCF2L) is able to drive lysosome clustering, a process that can antagonize lysosome transport by kinesin motors in heterologous cells.

### MCF2L drives dynamic clustering of lysosomes at AIS via reversible dimerization

Considering its enrichment at the AIS of neurons and its clustering activity, we then investigated whether MCF2L also modulates lysosome organization at AIS. In cultured hippocampal neurons, overexpression of MCF2L-GFP induced Lamp1-mCherry clustering at AIS, forming thick and continuous patch-like structures that were distinct from the diffuse or punctate patterns observed in control neurons expressing GFP alone (Fig. 6A). SIM-based imaging of native neurons free from exogenous MCF2L expression further revealed that AIS (here defined as the proximal 0-50 μm axonal region from the soma) inherently harbors a significantly higher density of Lamp1-mCherry puncta compared to distal axonal segments (500-550 μm from the soma). Moreover, AIS-localized lysosomal puncta were also significantly larger than their distal counterparts (Fig. 6B, 6D). This spatial enrichment suggests an intrinsic lysosome clustering mechanism at AIS. Importantly, MCF2L overexpression further amplified this size disparity, yielding dramatically enlarged lysosomal assemblies at AIS relative to both the control neurons and the distal axons of MCF2L-expressing neurons (Fig. 6C, 6D). Collectively, these findings indicate that MCF2L exerts an AIS-localized role on lysosomal clustering.

**Fig 6.**
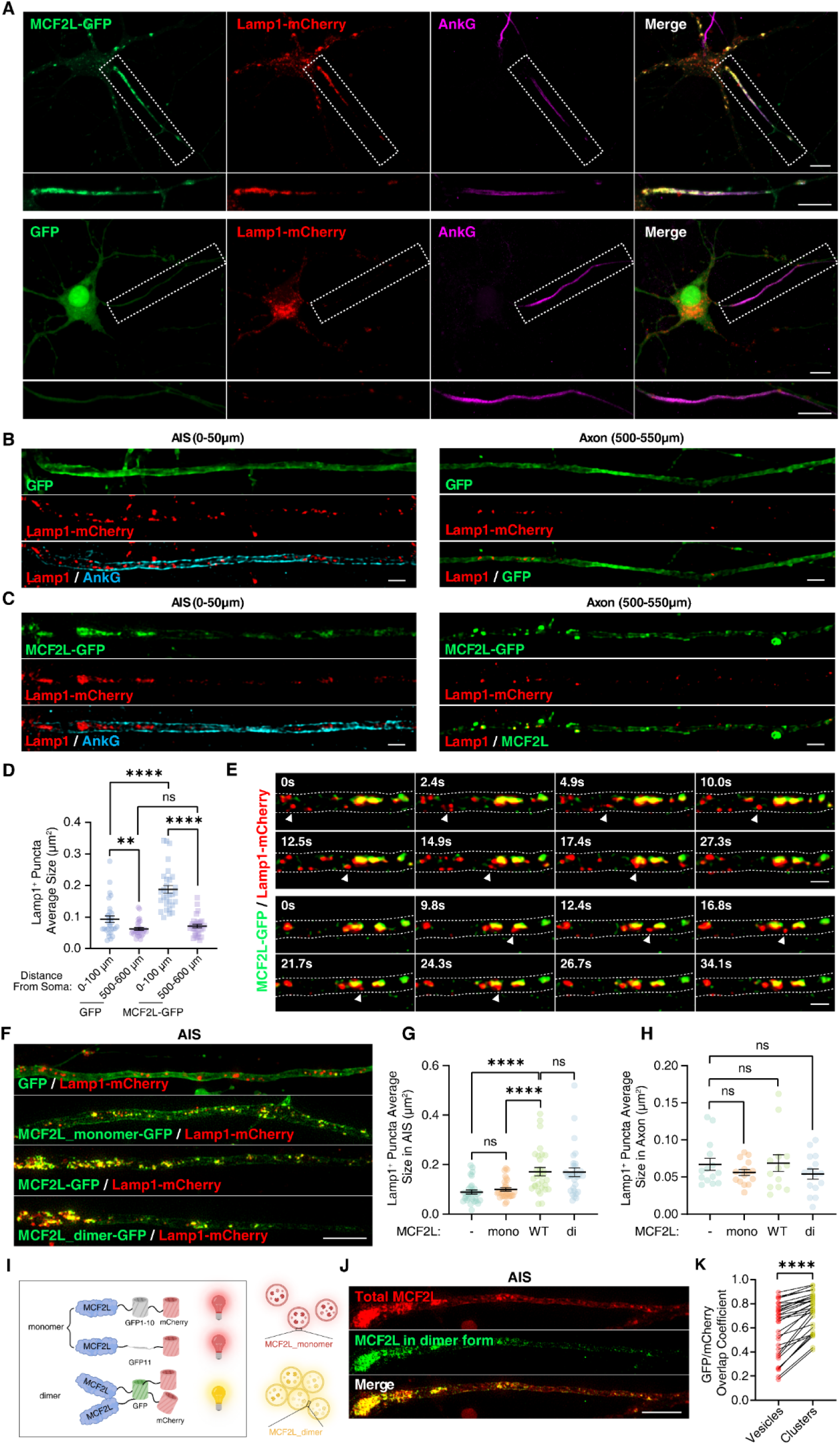
MCF2L preferentially drives dynamic clustering of lysosomes at the AIS via reversible dimerization. **A.** Representative images of neurons expressing GFP or MCF2L-GFP together with Lamp1-mCherry, and immunostained for AnkG. Scale bars, 10 µm. **B-C.** SIM images of neurons expressing GFP (**B**) or MCF2L-GFP (**C**) together with Lamp1-mCherry, and immunostained for AnkG. The AIS and distal axon were defined as 0-50 μm and 500-550 μm from the soma. Scale bars, 2 µm. **D.** Quantification of Lamp1 puncta size in different axonal regions. Mean ± SEM, three independent experiments; n=30 axons. **p=0.0088, ****p<0.0001; ns, not significant, two-tailed unpaired t test. **E.** Time-lapse SIM images of MCF2L-GFP and Lamp1-mCherry in the AIS, showing large MCF2L/Lamp1 clusters releasing and recapturing smaller Lamp1-positive vesicles. Scale bar, 1 μm. **F-H.** MCF2L-mediated lysosome clustering at AIS depends on its dimerization. **F.** SIM images of AIS in neurons expressing GFP, MCF2L-GFP, MCF2L_monomer-GFP, or MCF2L_dimer-GFP together with Lamp1-mCherry. Scale bar, 5 μm. **G-H.** Quantification of Lamp1-positive cluster sizes in AIS (**G**) and axon (**H**) of neurons expressing the indicated constructs. **G**, n=30 neurons. ****p<0.0001; ns, not significant, Mann-Whitney test. n=14 neurons for GFP, n=15 for MCF2L_monomer-GFP and MCF2L_dimer-GFP, and n=13 for MCF2L-GFP. ns, not significant, two-tailed unpaired t test. Data are shown as mean ± SEM from three independent experiments. **I-K.** MCF2L predominantly exists as the oligomerized form in AIS. **I.** Schematic diagram of the MCF2L-split-GFP chimera design. GFP fluorescence is reconstituted when MCF2L oligomerizes. **J.** SIM images of MCF2L-split GFP-mCherry expressed in the AIS. Scale bar, 5 μm. **K.** Quantification of GFP/mCherry overlap coefficient in individual vesicles or clusters within AIS. Mean ± SEM, three independent experiments; n=30 neurons. ****p<0.0001, Wilcoxon matched-pairs signed-rank test.

Live super-resolution imaging further delineated the dynamic behaviors of MCF2L- and Lamp1-labeled vesicles at AIS. We observed two distinct but interconvertible Ly/LE populations, classified as “clusters” (Lamp1-positive, >0.25 μm² area, >500 nm diameter) or “vesicles” (smaller) according to established size criteria ^86,87^. These two populations underwent continuous exchange. As shown in Fig. 6E, a small Lamp1-positive vesicle, indicated by an arrowhead, underwent rapid axonal transport, contacted and fused with an MCF2L-positive cluster, and became trapped (upper panel, and Movie 3). In another example, a small Lamp1-positive vesicle budded from a large cluster, underwent rapid axonal transport and was reincorporated into a distal cluster (lower panel and Movie 4). Interestingly, all large Lamp1-mCherry clusters also contained MCF2L-GFP, but small vesicles were frequently devoid of MCF2L-GFP signals or had only reduced signals (Fig. 6E), suggesting that local dissociation or reduction of MCF2L may accompany the formation of smaller and more mobile lysosomes in axons.

We then examined if the dimerization of MCF2L is important for lysosome clustering at AIS. WT, monomeric, and locked-dimer MCF2L variants all colocalized with Lamp1-positive puncta. However, the monomeric MCF2L mutant formed small and sparse puncta at AIS, whereas the locked-dimer mutant formed substantially larger AIS-localized puncta (Fig. 6F, 6G). Notably, in distal axonal regions, Lamp1 puncta size did not differ significantly among neurons expressing the WT, monomeric, or locked-dimer MCF2L variants (Fig. 6H). It is also noted that the increase in puncta size in AIS induced by the locked dimer is comparable to that induced by WT MCF2L. This could be explained by near saturation of the WT MCF2L dimer at AIS (likely by the total amount of gAnkG available in AIS) and thus it essentially behaves like a locked dimer. This interpretation fits with the very similar AIS clustering pattern and cluster size of WT MCF2L and the locked dimer as shown in Figure 6F.

We reasoned that enrichment of MCF2L by the high local concentration of gAnkG at AIS may promote MCF2L dimerization, thereby enhancing MCF2L-mediated lysosome clustering in this compartment. To verify this in situ, we employed a split-GFP reconstitution assay, in which MCF2L was fused separately to non-fluorescent GFP1-10 and GFP11 fragments. GFP fluorescence is reconstituted only when MCF2L molecules dimerize and bring the split GFP fragments into proximity (Fig. 6I). Upon co-transfection into neurons, bright punctate GFP signals were detected at AIS, whereas GFP fluorescence was sparse and rare in distal axonal regions (Fig. 6J). Moreover, large lysosomal clusters exhibited significantly higher split-GFP reconstitution frequency compared to small lysosome “vesicles” (Fig. 6K). Together, these results demonstrate that MCF2L undergoes reversible concentration-dependent dimerization and that this dimerization is required for dynamic lysosome clustering at AIS.

### MCF2L-mediated clustering and tethering to gAnkG cooperatively constrain lysosomal mobility at AIS

Live-cell tracking of Lamp1-positive vesicles in cultured neurons revealed that they move substantially more slowly within AIS than in distal axons, with average velocities of 0.58 ± 0.03 µm/s and 0.94 ± 0.05 µm/s, respectively (Fig. 7A-7C). This AIS-specific deceleration is consistent with previous observations of VAMP2-labeled vesicle dynamics in this compartment **^56^**, and suggests the existence of a site-specific mobility control mechanism restricting Ly/LE mobility at AIS. Further analysis of the relationship between Lamp1 puncta sizes and mobilities revealed that clustered Ly/LEs exhibited a marked reduction of mobile (with moving speed > 0.01 μm/s) fractions (Fig. 7D). Particularly, when the cluster size exceeds a defined size threshold (larger than the resolution limit of our confocal imaging), none of these clusters could undergo fast movements (Fig. 7D). These observations indicate that Ly/LE clustering is associated with their restricted trafficking.

**Fig 7.**
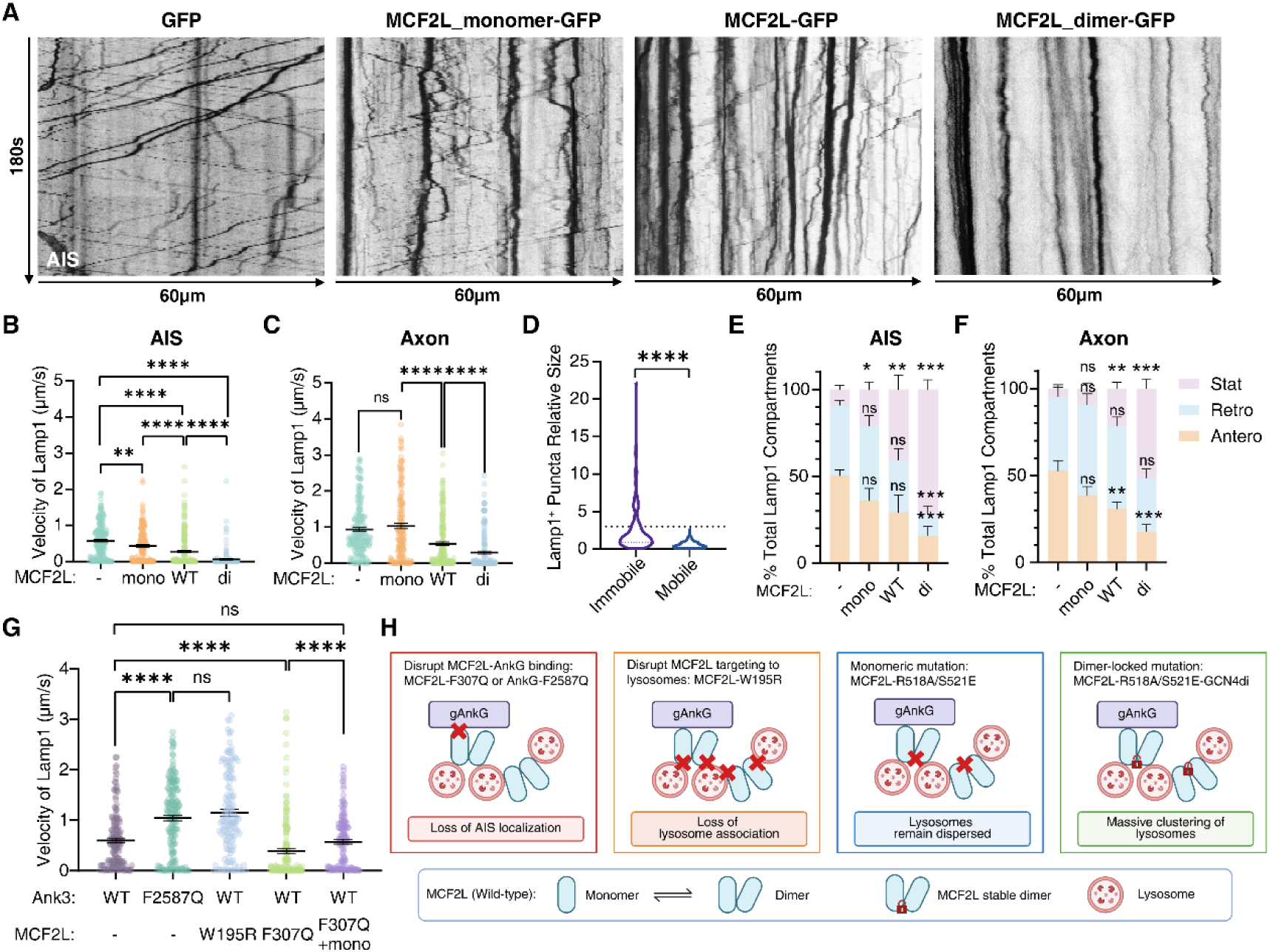
gAnkG-MCF2L-mediated clustering and tethering cooperatively restrict lysosomal transport at AIS. **A.** Representative kymographs of Lamp1 tracks in axons expressing GFP, MCF2L_monomer-GFP, MCF2L-GFP, or MCF2L_dimer-GFP. **B-C.** Quantification of Lamp1 transport speed in AIS (**B**) and axon (**C**) of neurons expressing the indicated constructs. Mean ± SEM, three independent experiments. **B**, n=206, 164, 185, and n=172 vesicles for GFP, MCF2L_monomer-GFP, MCF2L-GFP, and MCF2L_dimer-GFP, respectively, from three different batches of cultures. **p=0.0011 and ****p<0.0001. **C**, n=187, 171, 185, and 179 vesicles, respectively, from three batches of cultures. ****p<0.0001; ns, not significant, Mann-Whitney test. **D.** Quantification of Lamp1-positive immobile or mobile puncta size in AIS. Mean ± SEM, three independent experiments; n=342 for immobile clusters and n=190 for mobile vesicles. ****p<0.0001, Mann-Whitney test. The dashed black line indicates a threshold above which all clustered puncta are immobile. **E-F.** Quantification of Lamp1 transport direction in AIS (**E**) and proximal axon (**F**) of neurons expressing the indicated constructs. Mean ± SEM, three independent experiments. **E**, *p=0.046, **p=0.0023, and ***p=0.0004; **F**, **p<0.01, ***p<0.001; ns, not significant, Mann-Whitney test. **G.** Quantification of Lamp1 transport speed in AIS of *Ank3*^WT/WT^ neurons expressing GFP, MCF2L-W195R-GFP, MCF2L-F307Q-GFP, or MCF2L-F307Q_monomer-GFP, and in *Ank3*^F2587Q/F2587Q^ neurons without MCF2L transfection. Mean ± SEM, three independent experiments; n=144, 149, 173, 142, and 177 vesicles, respectively. ****p<0.0001; ns, not significant, Mann-Whitney test. **H.** Schematic diagram of the mutant constructs used in this study. MCF2L-F307Q and AnkG-F2587Q disrupt the MCF2L-AnkG interaction. The W195R mutation abolishes lysosomal targeting of MCF2L; the R518A/S521E mutation maintains MCF2L in a monomeric state; the R518A/S521E-GCN4di chimera locks MCF2L in a dimeric state.

Given the ability of MCF2L in inducing lysosomal clustering at AIS, we next investigated whether MCF2L, and its oligomerization state contribute to size-dependent regulation of lysosome mobility. MCF2L overexpression markedly reduced the average velocity of lysosome transport by increasing the immobile fraction in both AIS and distal axon, without altering the anterograde-to-retrograde transport ratio (Fig. 7A-7C and 7E, 7F). This inhibitory effect was markedly enhanced by expression of the locked-dimer MCF2L (see Fig. 7H for a summary of each variant), which rendered most AIS-localized Ly/LEs static (Fig. 7B, 7E). These data underscore that MCF2L-induced clustering inhibits lysosome transport in both the AIS and distal axon and that dynamic dimer-to-monomer conversion of MCF2L may facilitate lysosomal mobilization. Compared with GFP control, overexpression of MCF2L_monomer caused only a mild reduction in lysosome velocity at AIS (Fig. 7B) and had no detectable effect on distal axonal transport (Fig. 7C). This modest AIS-specific inhibition likely reflects the ability of monomeric MCF2L to tightly bind to gAnkG and tether lysosomes to the AIS scaffold, even in the absence of MCF2L-mediated clustering.

To determine whether the gAnkG-MCF2L-lysosome axis is essential for controlled lysosome transport at AIS, we employed neurons from *Ank3*^F2587Q/F2587Q^ mice. Lysosome transport at AIS was significantly faster in *Ank3*^F2587Q/F2587Q^ neurons (1.01 ± 0.06 µm/s) than in WT neurons (0.60 ± 0.05 µm/s) (Fig. 7G). Consistently, expression of the lysosome-targeting-deficient MCF2L mutant (W195R, see Fig. S4F), which retains gAnkG binding and AIS localization (Fig. S10A), similarly increased lysosome transport speed (Fig. 7G), likely by competitively inhibiting endogenous MCF2L binding to gAnkG. Conversely, overexpression of the gAnkG-binding-deficient MCF2L-F307Q mutant in WT neurons still impaired lysosome transport, consistent with its intact lysosome clustering capacity (Fig. 7G, 7H). Supporting this interpretation, expression of a monomeric MCF2L variant that also carried the F307Q mutation (i.e., F307Q, R518A and S521E triple mutation) had no detectable impact on lysosome transport velocity (Fig. 7G, 7H). These results distinguish two separable activities of MCF2L at AIS: gAnkG-dependent tethering and dimerization-dependent clustering of lysosomes.

Taken together, the above findings elucidate a dual regulatory mechanism for AIS lysosome mobility (Fig. 7H). High-affinity binding of MCF2L to gAnkG enriches MCF2L at AIS, where its high local concentration favors MCF2L dimerization and lysosomal clustering, thereby reducing lysosome mobility. Concurrently, direct binding of MCF2L-tethered lysosomes to gAnkG, which is highly concentrated at AIS and organizes the stable periodic membrane-cytoskeletal superstructures of AIS **^88^**, further impedes lysosome movement. The dynamic monomer-dimer equilibrium of MCF2L may allow small lysosomal vesicles to dissociate from larger clusters and resume axonal transport, although the upstream signal(s) governing this release remain to be defined.

### AIS as a potential quality-control checkpoint to trap impaired lysosome clusters

AIS is known to act as a selective barrier that prevents somatodendritic-specific vesicles from entering the axon **^56,89,90,58,78^**. However, whether AIS differentially regulates organelle transport based on subpopulation identity or functional state remains unclear. Given that MCF2L mediates lysosome clustering and transport inhibition at AIS, we asked whether the two distinct subpopulations, clustered and non-clustered lysosomes, differ in function (e.g., acidity). We employed a pH-sensitive reporter pHLARE (pH Lysosomal Activity REporter) **^91^**, which combines sfGFP-Lamp1 with pH-insensitive mCherry. In this system, acidic lysosomes quench sfGFP fluorescence, whereas less acidic compartments retain sfGFP signal. “vesicles” exhibited markedly reduced sfGFP-Lamp1 signals relative to “clusters” (Fig. 8A, 8B), meaning that MCF2L-associated lysosomal clusters retained at AIS are less acidic than their fast-moving, non-clustered counterparts. These high luminal pH Lamp1-positive clusters could correspond to multiple endolysosomal intermediates (e.g., late endosomes, amphisomes or incompletely acidified lysosomal hybrids) or impaired lysosomes.

**Fig 8.**
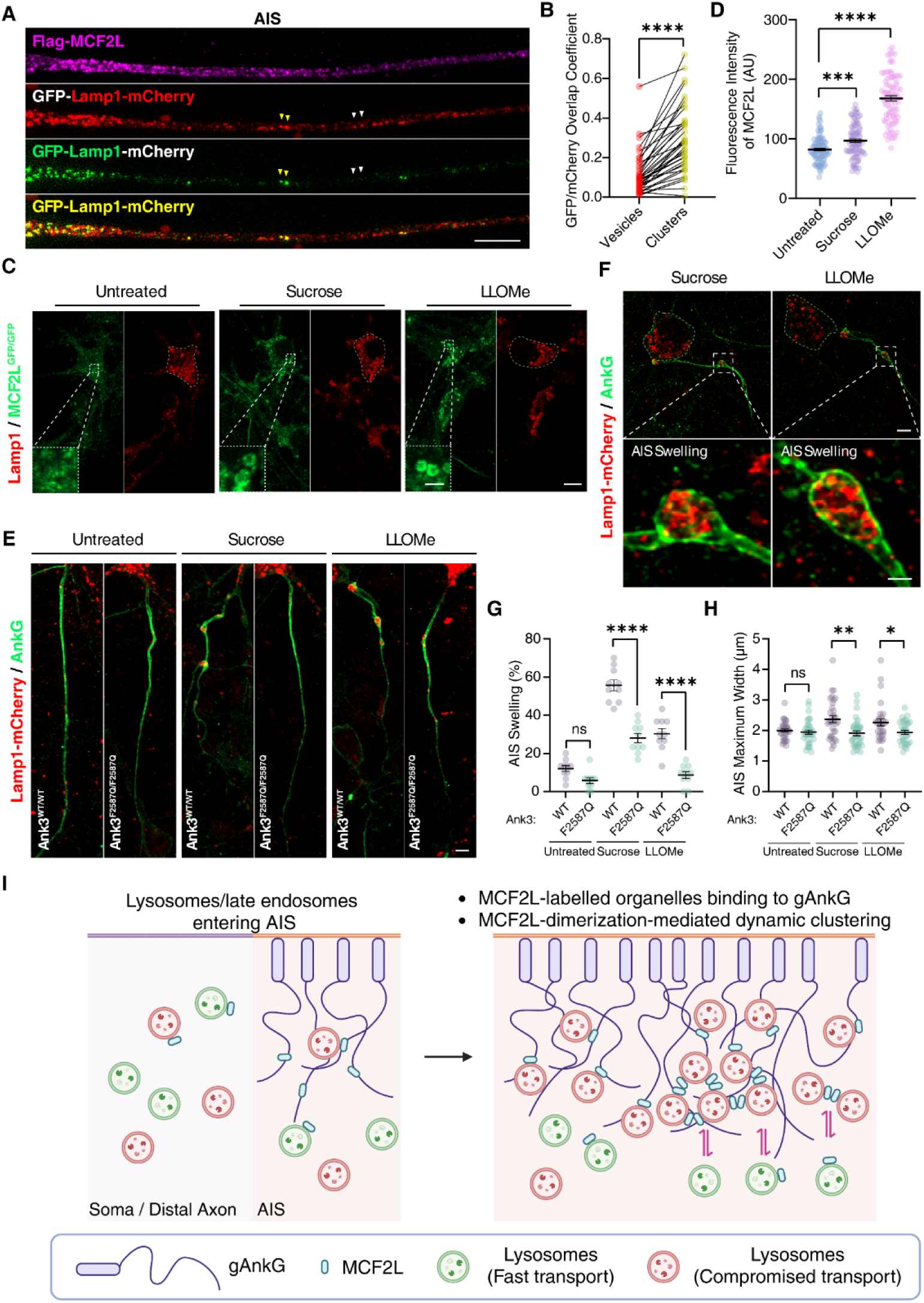
AIS-clustered lysosomes exhibit reduced acidity and integrity. A-B. Clustered lysosomes at AIS have reduced acidity. **A.** SIM image of AIS from a neuron expressing Flag-MCF2L and GFP-Lamp1-mCherry, pHLARE. Yellow arrows indicate GFP/mCherry-positive lysosomes with less acidic lumens. Red arrows indicate acidic lysosomes with quenched GFP signals. Scale bar, 5 μm. **B.** Overlap coefficient of the GFP-Lamp1/Lamp1-mCherry in individual lysosomal vesicles or clusters within AIS. Mean ± SEM, three independent experiments; n=37 neurons. ****p<0.0001, Wilcoxon matched-pairs signed-rank test. **C-D.** Sucrose or LLOMe treatment increases endogenous MCF2L protein intensity in neuronal lysosomes. **C.** Representative SIM images of MCF2L^GFP/GFP^ neurons under untreated, sucrose-treated or LLOMe-treated conditions stained for endogenous Lamp1. Scale bar, 10 µm. Lower panel, Dashed lines indicate manifested regions, scale bar, 2 µm. **D.** Quantification of average MCF2L fluorescence intensities on the Lamp1-positive vesicle membranes under the indicated conditions. Mean ± SEM, three independent experiments; n=120 vesicles, ****p<0.0001, Mann-Whitney test. **E-H.** Swollen or damaged lysosomes are preferentially retained at AIS of *Ank3*^WT/WT^ neurons. **E.** Representative confocal images of Lamp1-mCherry AAV-infected *Ank3*^WT/WT^ or *Ank3*^F2587Q/F2587Q^ neurons treated with sucrose or LLOMe and immunostained for AnkG. Scale bar, 5 μm. **F.** Representative SIM images showing the AIS swelling in sucrose- or LLOMe-treated neurons. Scale bars, 5 μm; magnified inset, 1 μm. **G.** Percentage of neurons with AIS swelling among untreated, sucrose-treated, and LLOMe-treated *Ank3*^WT/WT^ or *Ank3*^F2587Q/F2587Q^ neurons. Mean ± SEM, three independent experiments; n=10 groups of 300 neurons. ****p<0.0001, two-tailed unpaired t test. **H.** Maximum AIS width in untreated, sucrose-treated, and LLOMe-treated *Ank3*^WT/WT^ or *Ank3*^F2587Q/F2587Q^ neurons. Mean ± SEM, three independent experiments; n=33 neurons. ns, not significant; *p=0.0297, Mann-Whitney test; **p=0.0012, two-tailed unpaired t test. **I.** Schematic model of MCF2L-regulated lysosome clustering/transport at AIS. (Left) Ly/LEs entering AIS are tethered by gAnkG through MCF2L. (Right) MCF2L also induces dynamic clustering of Ly/LEs, reducing their motility. Small Ly/LE vesicles released from large clusters become mobile and transport competent.

Lysosomal damage, characterized by compromised membrane integrity or dysregulation of luminal acidity, is a pervasive physiological event that occurs continuously in living systems, and is particularly prevalent during aging, diseases and environmental stresses **^24,92,93^**. In neurons, stringent quality-control mechanisms are essential to prevent these potentially toxic organelles from accessing the vulnerable axonal compartment, but knowledge of this process is limited. We therefor asked whether MCF2L is linked to this process. We induced two distinct forms of lysosomal stress/damage in MCF2L^GFP/GFP^ neurons: sucrose treatment, which causes osmotic swelling, and L-leucyl-L-leucine methyl ester (LLOMe) treatment, which induces lysosomal swelling and membrane rupture **^67,94^**. Interestingly, following induction of lysosomal stress or damage by sucrose or LLOMe, endogenous GFP-tagged MCF2L became strikingly enriched on swollen lysosomal membranes marked by endogenous Lamp1 (Fig. 8C, 8D), suggesting that MCF2L can respond to stress-associated changes in lysosomes and may facilitate their retention at AIS.

We further tested whether the gAnkG-MCF2L interaction is required for the retention of stressed/damaged lysosomes at AIS. WT and *Ank3*^F2587Q/F2587Q^ neurons were treated with sucrose or LLOMe. Under both conditions, WT neurons developed pronounced axonal swellings at AIS (Fig. 8E, S10B). These swelling bumps were filled with large clusters of enlarged lysosomes that had lost their transport capability (Fig. 8F), indicating that stressed or damaged lysosomes are retained at AIS. Compared to untreated controls, WT neurons demonstrated a marked increase in the proportion of AIS population with lysosome-enriched swellings (Fig. 8G) and in the maximal width of these swellings (Fig. 8H). In striking contrast, sucrose- or LLOMe-induced AIS swellings were substantially less frequent and less pronounced in *Ank3*^F2587Q/F2587Q^ neurons (Fig. 8G, 8H), suggesting that the gAnkG-MCF2L interaction is required for efficient trapping of stressed or damaged lysosomes at AIS.

Collectively, our findings suggest a model for lysosomal quality control at AIS (Fig. 8I). Central to this model is the gAnkG-MCF2L axis, which suppresses lysosomal transport through two complementary mechanisms: it directly tethers lysosomes to AIS, thereby slowing their fast axonal transport; and it elevates local MCF2L concentration, promoting its oligomerization and consequently driving lysosomal clustering. MCF2L-clustered Lamp1-positive compartments are less mobile and are preferentially associated with less acidic or damaged lysosomes. Thus, the gAnkG-MCF2L complex may act as a sensor for monitoring the quality or maturation states of endolysosomal vesicles before they pass through AIS. Functionally competent lysosomes may dissociate from this AIS-associated quality-control gate and resume fast axonal transport, although the signals that regulate clustering of less mature or impaired lysosomes at AIS and the release of lysosomes from gAnkG-MCF2L trapped clusters remain to be identified. This AIS-mediated lysosome quality-controlling mechanism will have deep implications for axonal homeostasis *in vivo*, particularly under aging or pathological conditions.

## Discussion

### The MCF2L-gAnkG interaction is an evolutionary gain for AIS-specific functions

Although AnkG has multiple splice isoforms in neurons, the 480 kDa gAnkG isoform is exclusively clustered at AIS and nodes of Ranvier, which are homologous structures of AIS. In this study, we show that MCF2L, a protein preferentially expressed in the nervous system, selectively binds the 480 kDa gAnkG isoform rather than other AnkG isoforms. This specificity is further supported by the observation that giant AnkB, a paralog of gAnkG that is distributed throughout axons but excluded from AIS ^95–97^, lacks the MCF2L-binding sequence despite sharing many other interacting partners with gAnkG. These findings suggest that the gAnkG-MCF2L interaction has been evolutionarily refined for AIS-specific functions. The interaction between MCF2L and gAnkG is highly specific and strong (K_d_ ∼17 nM). Notably, the corresponding binding sequences in the two proteins emerged concurrently and remain highly conserved during the emergence of myelinated axons in cartilaginous fishes, a significant milestone in nervous system evolution that enabled rapid electric signal conduction over long distances ^98–101^. Long myelinated axons impose substantial homeostatic challenges, including high metabolic demand, increased vulnerability to oxidative stress and transport disruption, structural susceptibility to mechanical stress, etc. Thus, the gAnkG-MCF2L-lysosome axis may represent a dedicated molecular player that evolved to safeguard axonal structural and functional integrity throughout the lifetimes of neurons in the brain. The specific and high affinity interaction between MCF2L and gAnkG (Fig. 1 and Fig. 2) and very concentrated enrichment of gAnkG at AIS provide a molecular mechanism explaining why MCF2L-mediated lysosome clustering occurs preferentially at AIS, rather than in distal axons.

### Investigating gAnkG-mediated AIS lysosome gating without perturbing the AIS structural integrity

Genetic studies have linked AnkG (*Ank3*) and other AIS proteins to neurodegenerative disorders, but the underlying mechanisms remain poorly understood ^49,78,102^. Selective vesicle gating at AIS has long been postulated to be critical for axonal homeostasis ^103^, yet direct *in vivo* evidence has been difficult to obtain. A major obstacle is that AIS gating relies heavily on gAnkG, the giant master scaffold protein essential for the overall structural integrity of AIS as well as for anchoring numerous AIS-resident proteins including ion channels, cell adhesion proteins, and various regulatory proteins ^78^. Deletion or depletion of gAnkG will inevitably lead to overall disruption of the AIS structure and function, making mechanistic dissection difficult or even impossible. Besides, the huge size of gAnkG and its neuron specific acquisition of an extremely long intrinsically disordered sequence of ∼2,600 aa also pose challenges in identifying molecular compositions within the scaffold protein for each specific function. To overcome such challenges, we identified a single point mutation of gAnkG (gAnkG_F2587Q) that specifically impairs active lysosome gating at AIS without compromising the AIS structural integrity. The gAnkG_F2587Q mutation allowed us to investigate how gAnkG at AIS controls lysosome quality via its binding protein MCF2L. By crossing the *Ank3*^F2587Q/F2587Q^ mouse mutant line with a Parkinson’s disease model (hSNCA-A53T) mice, we provide *in vivo* evidence showing that impairment of the AIS-gated lysosomal trafficking exacerbates disease progression in the mouse model. It will be interesting to investigate in the future whether gAnkG may also regulate other cellular organelle quality-control and/or trafficking in axons via binding to certain organelle-specific proteins.

### Subcellular organelle clustering

Clustering has been observed for multiple distinct organelles, including synaptic vesicles, endosomes, lipid droplets, melanosomes, mitochondria and lysosomes, and often carries functional implications. Synaptic vesicle clusters serve as storage pools that support rapid neurotransmitter release upon depolarization ^104^, and thus influence overall synaptic functions ^104–106^. Similarly, endosome clustering facilitates efficient cargo sorting, exchange, and signaling by juxtaposing adjacent endosomes and promoting coordinated vesicle trafficking and maturation ^107,108^. Lipid droplet clustering facilitates their fusion and can modulate metabolic coupling with mitochondria during nutrient stress, thereby influencing cancer cell survival and tumor progression ^109,110^. Melanosome clustering in the perinuclear region can function as a non-degradative reservoir or protective structure that preserves melanosome integrity and function, contributing to photoprotection against ultraviolet radiation ^111–113^. Intriguingly, mitochondria also exhibit clustering behavior, particularly near the soma-facing entrance of AIS, where relatively immobile mitochondrial clusters may contribute to the maintenance of axonal Tau protein sorting ^114^ and calcium regulation ^115^. Although less studied in neurons, perinuclear lysosome clustering in non-neuronal cells coordinates with lysosome distribution, maturation and mTOR activity, particularly during metabolic stresses ^25,116^. Our findings further extend this principle: lysosome clustering mediated by lysosomal membrane-associated proteins, such as MCF2L, restricts kinesin-driven lysosomal transport. In non-neuronal cells, this process contributes to the regulation of lysosomal positioning and distribution within the cell. In neurons, such subcellular compartment-specific lysosome clustering is coupled to lysosome motility and quality control. More broadly, vesicle and organelle clustering may represent a common cellular strategy to coordinate organelle biogenesis, distribution, damage repair, and function under both physiological and pathophysiological conditions.

### Limitations and future directions

Several important questions regarding MCF2L-mediated lysosome clustering at the neuronal AIS remain unresolved. For example, how lysosomes are processed and subsequently released from the MCF2L-mediated clusters is not known, and it is unclear whether AIS clustering is a specific stress response, a sorting surveillance mechanism, or a routine trafficking process of lysosomes. The α-synuclein-A53T mouse model data present in Figure 3 of this study hint that the MCF2L-gAnkG axis may be a stress-response mechanism. However, this hypothesis requires rigorous future testing.

Previous studies indicate that most mature lysosomes remain in the soma, whereas a mixture of predominantly immature and some mature lysosomes enters the axon, with immature lysosomes maturing during retrograde transport to the soma ^19^. In our study, the Lamp1-positive vesicles clustered and retained by MCF2L-gAnkG at AIS were less acidic than their non-clustered counterparts (Fig. 8A-B). This finding should not be simply taken to mean that immature lysosomes are selectively trapped at AIS while mature lysosomes are released under normal cellular conditions. It is possible that AIS, via the MCF2L-gAnkG axis, acts as a surveillance apparatus monitoring lysosomal states. Under steady-state conditions, lysosomes in axons generally undergo retrograde transport, following the maturation-along-transport mode described in the literature. It is possible that, upon stress, AIS may impose another layer of control over lysosome trafficking in neuronal axons, limiting impaired lysosomes from entering and accumulating along long, slender axons.

Additionally, the properties of clustered lysosomes in AIS will need to be further characterized. In the current study, we mainly used Lamp1 to mark AIS lysosomes. In neurons, Lamp1 is present on a heterogeneous population of endolysosomal organelles, many of which are non-degradative ^117–119^. The less acidic Lamp1-positive compartments may include immature or incompletely acidified endolysosomes, amphisomes or autolysosomes, impaired lysosomes, lysosome-related organelles, or Lamp1-bearing biosynthetic and trafficking intermediates ^120^. Conversely, more acidic, non-clustered vesicles are not necessarily fully mature lysosomes. Further characterization of these Lamp1-positive vesicular organelles is required.

Regarding MCF2L-mediated dynamic lysosome clustering, multiple molecular processes are possible. Mildly damaged lysosomes may undergo local repair and maturation, whereas irreversibly damaged organelles may be transported retrogradely to the soma or eliminated by local lysophagy. Functionally competent lysosomes are released from AIS-associated clusters to resume fast axonal transport. The release of lysosomes from AIS may involve dissociation of MCF2L from lysosomal membranes or a reduction of MCF2L density on the lysosome surface, either events could trigger lysosome de-clustering. Consistent with this possibility, our SIM imaging showed that small, rapidly moving lysosomes released from large clusters contained reduced or even undetectable MCF2L fluorescence signals (Fig. 6E, Movie 3 and Movie 4). Alternatively, neuronal activity-dependent dissociation of the MCF2L dimer or disruption of the MCF2L-gAnkG interaction may lead to lysosome de-clustering and their release from the AIS. Addressing these questions in the future may clarify how AIS actively regulates lysosome trafficking, repair, maturation, and disposal in neuronal axons.

## Methods

### Plasmid constructs

The full-length cDNA encoding human MCF2L (NCBI sequence ID: NM_024979.4) was purchased from YouBio Inc., China. Plasmids encoding the 270-kDa and 480-kDa rat *Ank3* isoforms (NCBI sequence ID: XM_039098838.2 and XM_039098834.2), human Lamp1 (NCBI sequence ID: NM_005561.4), human COX8A/mito (NCBI sequence ID: NM_004074.3), human Rab7A (NCBI sequence ID: NM_004637.6), human TfR (NCBI sequence ID: NM_001128148.3), and human TGOLN2/TGN38 (NCBI sequence ID: NM_006464.4) were gifts from Dr. Vann Bennett (Duke University, USA). pCAGGS-mTagBFP2-T2A-iCre (v2) (#125822), pFX-mCherry-EEA1 (#174452), Lamp1-iRFP (#136570), and GFP-Lamp1-mCherry (#164477) constructs were purchased from Addgene. The coding sequence of mouse Arl8b (NCBI sequence ID: NM_026011.3) was amplified from a mouse cDNA library. The coding sequence of FRB and FKBP were amplified from PM-FRB-mRFP-T2A-FKBP-5-ptase plasmid (Addgene, #40896). The coding sequence of GFP1-10 and GFP11 were amplified from pcDNA3.1-GFP(1-10) (Addgene, #70219) and pEGFP-GFP11-Clathrin light chain (Addgene, #70217), respectively. The coding sequences of SNCA and Rab5A were amplified from pCMV-SNCA(human)-3×FLAG-Neo (miaolingbio, #P42768) and pCMV-mCherry-RAB5A(human)-Neo (miaolingbio, #P41491), respectively. All truncations and mutations used in this study were generated by the standard PCR-based methods. PCR products were inserted into modified pET32a vector (pET.32M.3C) for bacterial expression of Trx-His_6_-tagged or Trx-3×StrepII-His_6_-tagged fusion proteins. For mammalian expression, fragments were cloned into pCAG-GFP (a gift from Dr. Gary Banker), pCAG-mCherry, pcDNA3.0-3×Flag or pmiRFP670-2 vectors. All constructs were verified by DNA sequencing.

### Animals and Ethics

All mice were housed in individually ventilated cages (a maximum of six mice per cage) within a specific-pathogen-free barrier facility at Shenzhen Bay Laboratory. The mice were maintained on a 12-h light/12-h dark cycle, at a consistent ambient temperature (21-23℃) and humidity (50-60%), with sterile pellet food and water ad libitum. All animal experiments were performed in accordance with the ethical guidelines of the Shenzhen Bay Laboratory animal experimentation committee.

### Ank3^F2587Q/F2587Q^ mice

The AnkG-F2587Q point mutation knock-in mouse line was obtained by CRISPR/Cas9-mediated genome engineering in C57BL/6J background (Cyagen Biosciences). Cas9-guide RNA (gRNA) target sequences were designed to the exon 37 to promote DNA cleavage and homology-directed repair. Silent mutations (AGG to CGC, AAA to AAG and, GGC to GGA) were introduced to prevent gRNA rebinding and recutting after repair. Heterozygous mice were backcrossed to WT C57BL/6J mice for five generations and then intercrossed to establish a homozygous line. The knock-in mice were born at approximately Mendelian ratios and reproduced normally. Mice were genotyped by PCR using tail genomic DNA. Primers used for genotyping were as follows: Primers A, 5’-TGAAGTGTCGCAGTTTTT-3’ and 5’-CTTAACCCATTCATCGTTGG-3’; Primers B, 5’-CAGAGTTACAGGGGTCTGAT-3’ and 5’-TTCTCAGTTTTGTCACGTTG-3’.

### hSNCA-A53T mice

hSNCA-A53T mice were obtained from GemPharmatech, Jiangsu, China (Strain NO. T054329). This strain was produced by introducing the human Synuclein A53T mutant under the control of the Thy promoter in the C57BL/6J background. Primers used for genotyping were as follows: Primers C, 5’-GCTGAGGTATTCATCATGTGCTCC-3’ and 5’-TGTCTTCTGGGCTACTGCTGTCA-3’; Primers D, 5’-TGCCTTCTGAGGAAGGGTATCAAG-3’ and 5’-TTTCAGGCATCTGGCTTGGTT-3’.

### Yeast two-hybrid screening

A commercial ULTImate yeast two-hybrid (Y2H) screen was performed by Hybrigenics Services (Evry-Courcouronnes, France). The bait construct contained amino acids 2329-2691 of rat AnkG cloned into the pB27 vector to generate an N-LexA-bait-C fusion. The bait was screened against a mouse adult brain cDNA library (Mouse Adult Brain_RP1). Two pB27 screen datasets, pB27_A and pB27_B, were generated using 20 mM and 10 mM 3-amino-1,2,4-triazole (3-AT), respectively, to reduce background growth. Prey clones were identified by sequencing the available 5’ and/or 3’ ends of the cDNA inserts and comparing the sequences with reference databases. The orientation, reading frame, and sequence identity of each prey fragment were determined during the bioinformatic analysis. Interaction confidence was evaluated using the Hybrigenics Global Predicted Biological Score (PBS), which ranks interactions according to the independent recovery of prey fragments and identifies highly connected prey domains or experimentally defined technical artifacts. For interaction-domain analysis, the selected interaction domain (SID) was defined as the amino acid sequence shared by prey fragments corresponding to the same reference protein. Known structural and functional domains were annotated using PFAM, SMART, TMHMM, Ncoils, and SignalP analyses.

### Expression and purification of recombinant proteins

Plasmids were transformed and expressed in BL21 (DE3) codon-plus *E.coli* at 16 °C for 16-20 h after induction by IPTG. The N-terminal Thioredoxin-His6-tagged (Trx-His6) or Trx-3×StrepII-His6-tagged proteins were purified using an Ni^2+^-NTA agarose affinity column followed by size-exclusion chromatography (SEC) in the final buffer containing 50 mM Tris-HCl, pH 7.8, 100 mM NaCl, 1 mM DTT, and 1 mM EDTA. When required, Trx-His_6_-tag was cleaved by HRV-3C protease at 4 °C overnight and removed by another step of SEC.

### Isothermal titration calorimetry (ITC)

The interactions between MCF2L_Spec1 and gAnkG_peptide were assessed by ITC using a MicroCal VP-ITC calorimeter (Malvern) at 25℃ in the assay buffer (50 mM Tris-HCl, pH 7.5, 100 mM NaCl, 1 mM EDTA, and 1 mM DTT). MCF2L_Spec1 (20 μM) and gAnkG_peptide (200 μM) were loaded into the sample cell and syringe, respectively. Each titration point contained an injection of 10 μL aliquot delivered with a 120-s interval. Data were analyzed with a one-site binding model in Origin 7.0 software.

### Size exclusion chromatography with multi-angle light scattering (SEC-MALS)

Protein oligomeric state and molecular weight were determined by SEC-MALS using an Agilent InfinityLab system with a miniDawn static light scattering and an Optilab differential refractive index detector (Wyatt). Samples (concentrations specified in the figure legends) were filtered (0.22 μm) and injected onto a Superose 12 10/300 GL column (GE Healthcare) equilibrated in 50 mM Tris-HCl, pH 7.5, 100 mM NaCl, 1 mM EDTA, and 1 mM DTT. Data were analyzed using ASTRA6 software (Wyatt).

### Circular dichroism spectroscopy

Circular dichroism (CD) spectra of purified tag-free MCF2L_Spec1 protein were acquired on a JASCO J-810 CD spectropolarimeter at room temperature using a 1-mm path-length cuvette. Spectra were collected over a wavelength range of 200-300 nm, and each spectrum represents the average of two repeated measurements. Protein samples were prepared at 20 μM in 25 mM Tris-HCl, pH 7.5, containing 50 mM NaCl, 0.5 mM EDTA and 0.5 mM dithiothreitol. For urea-induced denaturation assays, spectra were collected in the same buffer supplemented with 1, 2, 2.5, 3, 3.25, 3.5, 4, 5, 6, or 7 M urea. All proteins used for CD analysis were purified after removal of affinity tags.

### Crystallization and structure determination

To determine the structure of the AnkG 2553–2604 / MCF2L_Spec1 complex, we engineered a fusion construct in which two spectrin repeats derived from human β2 spectrin (residues 1811-2015) were fused to the C-terminus of MCF2L_Spec1 (residues 196-326). This design helps to expand the crystal packing interface, thereby facilitating crystallization. AnkG 2553-2604/MCF2L_Spec1-β2 spectrin crystals were obtained by sitting-drop vapor diffusion at 16 ℃ in 4.0 M ammonium acetate and 0.1 M sodium acetate trihydrate (pH 4.0). Crystals were cryoprotected in 25% glycerol and flash-frozen in liquid nitrogen. X-ray diffraction data were collected at the Shanghai Synchrotron Radiation Facility (SSRF). Data were processed and scaled, and the structure was solved by molecular replacement using the β2 spectrin structure (PDB: 3EDV) as the search model, following the methodology described earlier ^121^. Iterative model building, refinement, and validation were performed using standard crystallographic software packages. Structural analysis and figure preparation were conducted using PyMOL. Statistics for the final model are summarized in Table S1.

### Immunoprecipitation and immunoblotting

MCF2L^GFP/GFP^ mouse brain tissue was homogenized in RIPA lysis buffer, centrifuged at 16,873 g for 10 minutes at 4℃, and the supernatant was diluted with 50 mM Tris-HCl, pH 7.5, 150 mM NaCl, and 1 mM EDTA. Lysates were incubated with anti-GFP nanobody agarose beads for 2 hours at 4℃ with gentle rotation. After washing, captured proteins were eluted with 2× SDS-PAGE loading dye and resolved by SDS-PAGE, transferred to PVDF membranes, and probed with mouse anti-GFP (Proteintech, #66002-1-Ig) antibody and secondary antibodies (Sigma-Aldrich, #A4416).

### Hippocampal neuron culture, transfection, immunostaining and chemical treatment

Primary hippocampal neuron cultures were prepared from postnatal 0∼1 day mouse brains. Hippocampi were dissected, dissociated, and plated on poly-L-lysine-coated coverslips (∼120,000 neurons/35-mm dish). Neurons were cultured in Neurobasal-A (Gibco, #12349015) with 10% fetal bovine serum (FBS, ThermoFisher, #10099141), 2% B-27 supplement (ThermoFisher, #17504044), and 1% GlutaMAX (ThermoFisher). After overnight incubation, the medium was replaced with Neurobasal-A containing 1% FBS, 2% B-27 supplement, and 1% GlutaMAX. Cultures were maintained at 37℃ in a humidified incubator with 5% CO2.

Hippocampal neurons at 6-7-days-*in vitro* (DIV) were transfected with Lipofectamine 2000 (ThermoFisher, #13778075) and subsequently fixed with 4% PFA containing 4% sucrose in PBS for 15 minutes at 37℃ before antibody staining. Fixed neurons were washed with PBS, permeabilized with 0.2% Triton X-100 for 10 min, and then blocked with 5% BSA in PBS for 1 hour. Primary antibodies were diluted in blocking buffer (5% BSA in PBS) and incubated at 4°C overnight. The following primary antibodies were used: rat anti-Lamp1 antibody (BD Biosciences, #553792), mouse anti-AnkG antibody (NeuroMab, #75-146), guinea pig anti-TRIM46 antibody (Synaptic Systems, #377308), mouse anti-Flag (Sigma-Aldrich, #F3165), and chicken anti-GFP (ThermoFisher, #A10262). The next day, neurons were washed with PBS and incubated with Alexa Fluor 594–conjugated secondary antibody (ThermoFisher, #A-21203, #A-11007), Alexa Fluor 647–conjugated secondary antibody (ThermoFisher, #A-21450, #A-31571), and Alexa Fluor 488–conjugated secondary antibody (ThermoFisher, #A78948) in blocking buffer for 1 hour at room temperature. Finally, the fixed neurons were imaged by a Zeiss LSM 880 laser-scanning confocal microscope or a Zeiss Elyra 7 at the Lattice SIM mode.

### Pharmacological induction of lysosome stress or damage in neurons

Cultured hippocampal neurons from MCF2L^GFP/GFP^ postnatal day 0∼1 mice were treated with either 100 mM sucrose for 24 hours or 1 mM LLOMe for 3 hours to induce lysosomal stress or damage. The treatments were initiated at DIV7 or DIV8, respectively. All neurons were fixed at DIV8 and immunostained for GFP and AnkG for imaging.

### Live-cell imaging of Lamp1 and quantification

Cultured hippocampal neurons were transfected with Lamp1-mCherry plasmids at DIV6 with Lipofectamine 2000. On DIV7, neurons were imaged on a Zeiss LSM 880 laser-scanning confocal microscope equipped with an AxioCam digital camera, a GaAsP detector, and a temperature- and CO_2_-controlled incubation chamber. The whole imaging stage and objectives were maintained at 37℃ and 5% CO_2_ in an enclosure. For kymograph generation and speed quantification, time-lapse images were acquired at AIS or mid-axon (∼500 μm away from the soma) with a Plan-Apochromat 40x/1.4 Oil DIC M27 objective, at a rate of 1 frame/s using the zoom and definite focus functions in the Zen acquisition and imaging software. Kymographs were obtained using the Kymograph Tracker plugin in Icy. Anterograde and retrograde motile vesicles were defined as particles showing a net displacement >3 μm in one particular direction, and stationary vesicles were defined as particles with a net speed <0.01 μm/s.

### Immunohistochemistry

Adult *Ank3*^F2587Q/WT^×hSNCA-A53T or *Ank3*^WT/WT^×hSNCA-A53T mice at desired time points were anesthetized via an intraperitoneal injection of an overdose of pentobarbital sodium. Upon confirmation of deep anesthesia (absence of pedal reflex), transcardial perfusion was initiated with 100 mL of ice-cold phosphate-buffered saline (PBS), followed by 100 mL of 4% paraformaldehyde (PFA) in PBS. Whole brains were carefully dissected and post-fixed in 4% PFA for 24 hours. Following fixation, tissues were dehydrated through a graded ethanol series, cleared in xylene, and embedded in paraffin blocks.

Coronal sections (5 µm thickness) were cut and mounted on glass slides. For immunohistochemical staining, sections were first deparaffinized in xylene and rehydrated through a graded alcohol series to PBS. Antigen retrieval was performed by incubating sections in 10 mM sodium citrate buffer (pH 6.0) and heating in an autoclave (5 min at high heat, followed by 15 min at low heat). Slides were then cooled to room temperature for 20 minutes. To minimize non-specific antibody binding, sections were incubated in a blocking buffer consisting of 10% (v/v) normal goat serum, 3% (w/v) fish gelatin, 1 mM EDTA, and 0.1% Tween-20 in PBS for 1 hour at room temperature.

Sections were subsequently incubated overnight at 4℃ with the following primary antibodies diluted in blocking buffer: anti-human α-synuclein (#ALX-804-258-L001, Enzo), and anti-phospho-α-synuclein (Ser129) (#23706S, Cell Signaling Technology). The following day, sections were washed thoroughly with PBS and incubated for 1-2 hours at room temperature with appropriate Alexa Fluor-conjugated secondary antibodies diluted in blocking buffer. Finally, sections were rinsed, briefly air-dried, and coverslipped using an antifade mounting medium containing DAPI for nuclear counterstaining (#P0131, Beyotime). Fluorescence images were acquired using a Zeiss LSM980 confocal laser-scanning microscope.

### Behavioral tests for *Ank3*^F2587Q/F2587Q^ mice

Behavioral assessments were performed in 2-month-old mice. Before each test, mice were allowed to habituate to the testing room for at least 1 h. Between trials, the apparatuses were thoroughly cleaned with 75% ethanol to minimize olfactory cues.

The open field test was used to assess anxiety-like behavior, exploratory activity, and locomotor function. Each mouse was placed individually in the center of a semi-transparent acrylic arena measuring 50 cm × 50 cm × 50 cm and allowed to explore freely for 10 min. Movement was recorded and analyzed using SMART video tracking software. The total distance traveled, time spent in the center area, and number of center entries were measured.

The elevated plus maze test was used to assess anxiety-like behavior. The apparatus consisted of two open arms (60 cm × 5 cm), two closed arms (15 cm × 5 cm, with 15-cm-high walls), and a central platform (5 cm × 5 cm), elevated 50 cm above the floor. Under controlled lighting conditions, each mouse was placed on the central platform facing an open arm and allowed to explore freely for 5 min. Time spent in the open arms and the number of open-arm entries were recorded and analyzed using SMART video tracking software.

The forced swimming test was used to assess immobility behavior. Mice were placed in a transparent cylindrical glass tank (50 cm high, 20 cm in diameter) containing water at a depth of 23 cm and a temperature of 25°C. Twenty-four hours before the formal test, mice underwent a 15-min training session in the water. On the test day, each mouse was placed in the tank for 6 min. Immobility during the final 4 min was recorded and analyzed using SMART video tracking software. Immobility was defined as floating without active struggling. Mice were towel-dried immediately after the test.

The tail suspension test was used to assess immobility behavior. Each mouse was suspended by the tail from a vertical metal rod using adhesive tape, with care taken to prevent contact with the floor or any other support. The test lasted 6 min, and immobility during the final 4 min was recorded and analyzed using SMART video tracking software.

### Behavioral tests for *Ank3*^WT/F2587Q^ × hSNCA-A53T mice

The rotarod test was used to assess motor learning and coordination. To acclimate to the rotarod apparatus (Panlab, Model: LE 8505, USA), mice were trained over three days with three daily trials at a constant speed of 5 and 10 rpm. During the acquisition session, mice underwent three trials, each separated by 45-s intervals. The latency to fall off the rod was recorded for each trial, with a maximum cutoff of 120 seconds, and the apparatus was cleaned with 75% ethanol between trials.

The pole test was used as a measure of motor coordination and function. Mice were placed head-down at the top of a vertical, gauze-wrapped wooden pole (50 cm in height, 1 cm in diameter) and the time to descend to the base was recorded. Preceding the actual evaluation, mice underwent a three-day training period (three test trials per day). On the test day, the mean latency to descend across three trials was calculated for each mouse.

The CatWalk Gait analysis was performed using the CatWalk XT system (version 10.5, Noldus Information Technology). Mice were first trained for one week to traverse the raised glass platform three times daily. For testing, animals voluntarily crossed the platform in a darkened room, and their footsteps were captured by a high-speed camera. The resulting videos were analyzed automatically by the CatWalk XT software to extract quantitative gait parameters.

In the tail suspension test, the hind limb suspension score was estimated during the first 10-15 seconds to elicit the clasping phenotype, which was scored on a scale from 0-4. 0: The hind limbs are normally spread. 1: One hind limb is not fully spread. 2: Inability to fully spread both hind limbs with normal toe extension. 3: The toes of both hind limbs are curled, and the thighs tremble involuntarily. 4: Fore and hind limbs gripped tightly with the body curled up.

### Statistical analysis

Statistical analyses and graph generation were performed using GraphPad Prism 10 (La Jolla, CA). Data are presented as mean ± standard error of the mean (SEM). Normality was assessed using the Shapiro-Wilk test. Two-group comparisons used an unpaired, two-tailed Student’s *t*-test (for normally distributed data), or the Mann Whitney test (for non-normally distributed data). Significance was set at p<0.05. Specific statistical tests, sample sizes (n), and P values are reported in figure legends.

## Acknowledgements

This work was supported by grants from the Major Program of Shenzhen Bay Laboratory (S201101002), the Fundamental and Interdisciplinary Breakthrough Plan of the Ministry of Education of China (JYB2025XDXM505), and the Key Research and Development Program of Guangdong Province (Grant No. 2023B0303010001) to M. Z. We thank the BL19U1 beamline at the National Facility for Protein Science Shanghai for X-ray beam time, Drs. Xiaochen Wang and Hong Zhang for their insightful comments of the manuscript.

## Author contributions

Y. L., K. C. and M. Z. conceptualized the project and designed the experiments. Y. L., K. C., Q. T., H. M., Y. H., and F. L. performed sample preparation and experiments. Z. L. solved the crystal structure. M. Z. and K. C. supervised the study. Y. L., K. C., R. Y. and M. Z. analyzed data. Y. L., K. C. and M. Z. wrote the original draft. All authors reviewed and edited the manuscript. M. Z. acquired the funding and coordinated the project.

