## Supplementary material for "Axon initial segment as an axonal lysosome trafficking control gate": Supplymentary figures and tables

Short title: *AIS as an axonal lysosome trafficking-control gate*

[#]These authors contributed equally to this work

### Figure S1

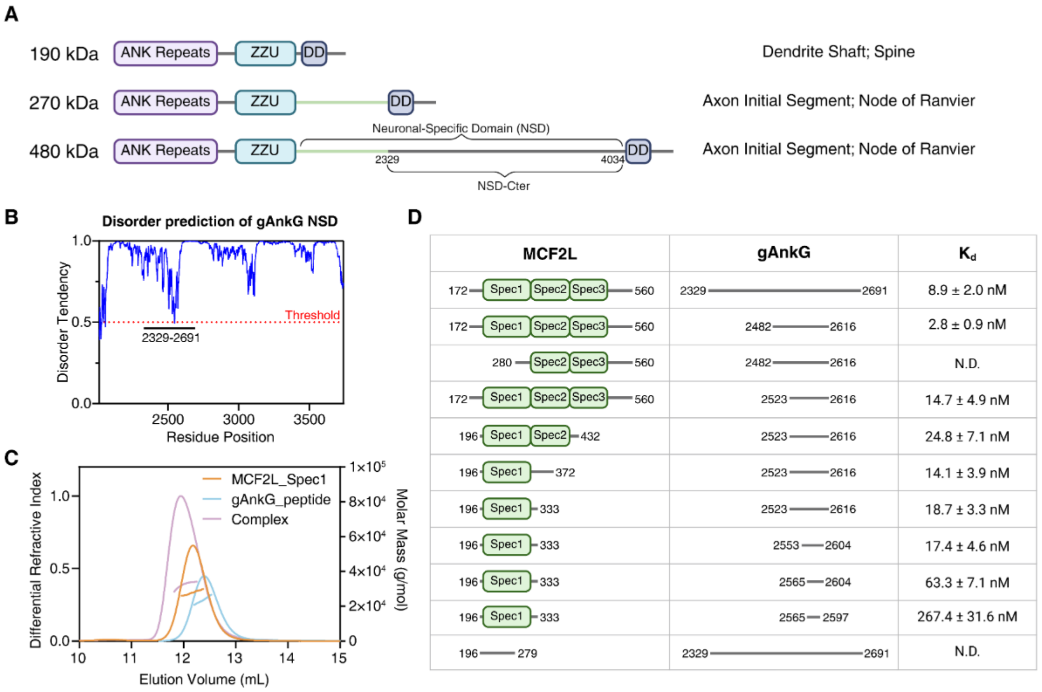


**Fig S1. Identification and characterization of the gAnkG-MCF2L interaction.**

**A.** Domain organization and subcellular localization of three major neuronal splice variants of AnkG. The 480 kDa variant contains a long intrinsically disordered insertion sequence (NSD-Cter), which is absent from the 270 kDa variant.

**B.** Disorder prediction of the entire gAnkG neuron-specific domain (NSD; aa 2014–3740) using IUPre2A (https://iupred2a.elte.hu/). A disorder probability score greater than 0.5 indicates intrinsically disorder.

**C.** SEC-MALS analysis showing stable complex formation between MCF2L_Spec1 and gAnkG_peptide.

**D.** Binding affinity guided mapping of key regions responsible for gAnkG-MCF2L interaction. Protein boundaries and corresponding dissociation constants measured by ITC are listed. N.D., not determined. The ITC binding curve for MCF2L Spec1 (aa 196-333) and gAnkG_peptide (aa 2553-2604) interaction pair (boxed in the panel) is shown in Fig. 1D.

### Figure S2

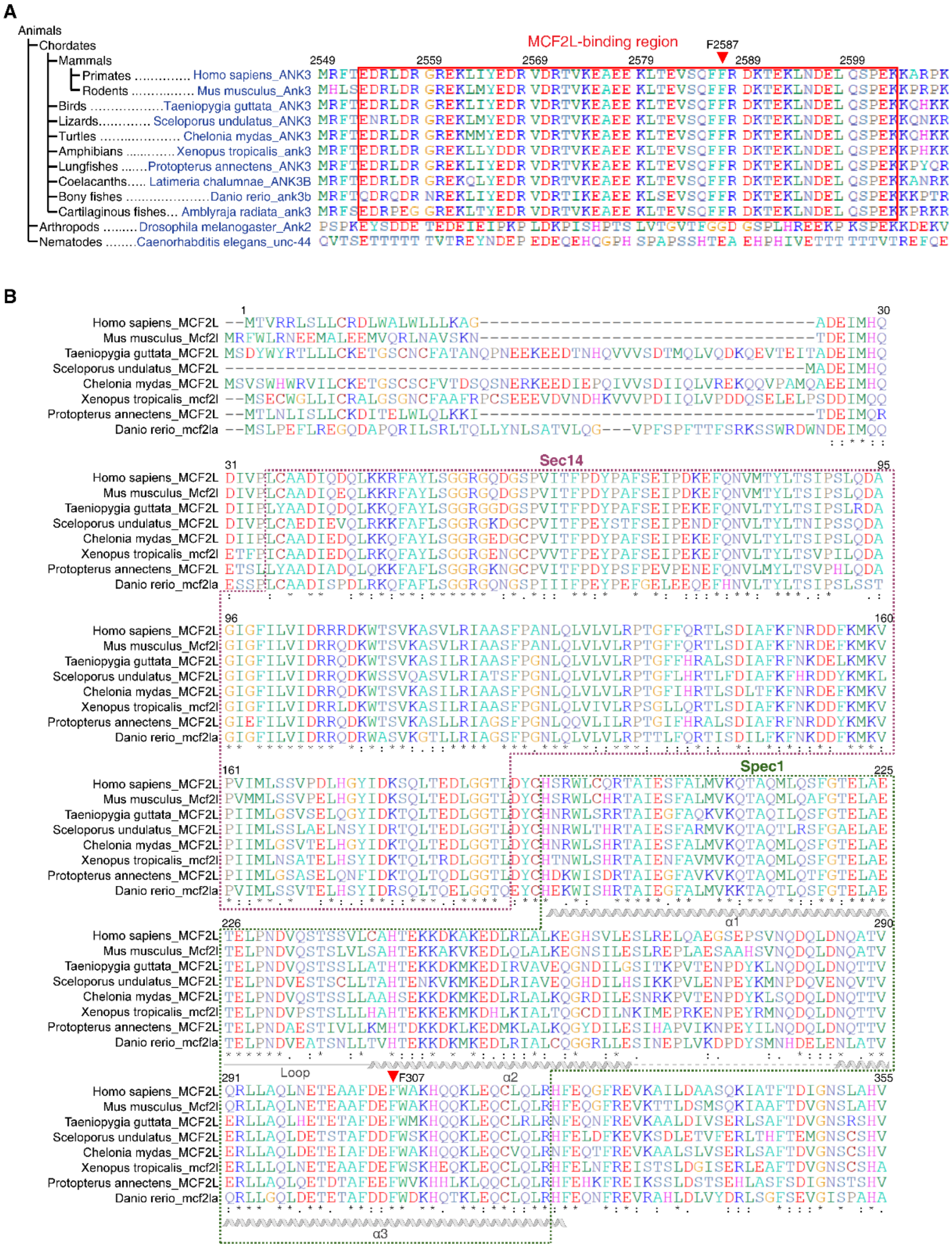

**Fig S2. Binding sequences in MCF2L and gAnkG are evolutionarily conserved.**

**A.** Evolutionary tree and sequence alignment of gAnkG_peptide segment (boxed in red) of gAnkG. The critical residue F2587 selected for mutagenesis is indicated by a red arrow. Note that *Stylophora pistillata* lacks giant ankyrins.

**B.** Sequence alignment of the MCF2L Sec14-Spec1 supramodule. The Sec14 is boxed in red and the gAnkG-binding region, Spec1, is boxed in green. The critical residue F307 selected for mutagenesis is indicated by a red arrow. Note that the sequence connecting the Sec14 and Spec1 domains are short and highly conserved.

#### Figure S3

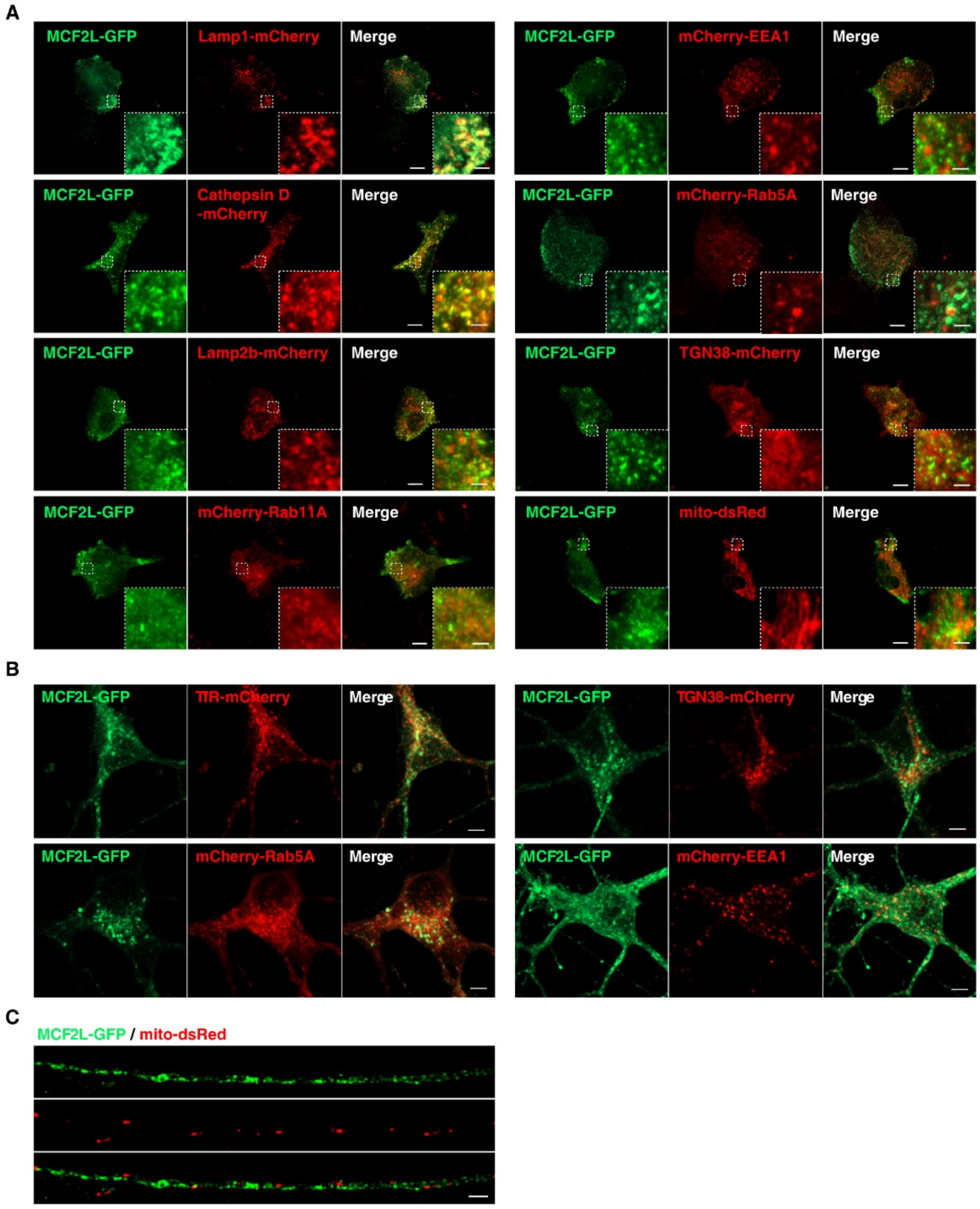


**Fig S3. MCF2L selectively colocalizes with lysosomes both in neurons and in non-neuronal cells.**

**A.** Representative confocal images of astrocytes expressing MCF2L-GFP together with various organelle markers, including Lamp1, Cathepsin D, Lamp2b, Rab11A, mito, EEA1, Rab5A, and TGN38. Scale bar, 10 μm; magnified inset, 2 μm.

**B.** Representative confocal images of cultured hippocampal neurons expressing MCF2L-GFP together with indicated vesicle markers, including TfR, TGN38, Rab5A, and EEA1. Scale bar, 5 μm.

**C.** SIM images showing that MCF2L-GFP does not colocalize with mitochondria in axon. Scale bar, 2 μm.

### Figure S4

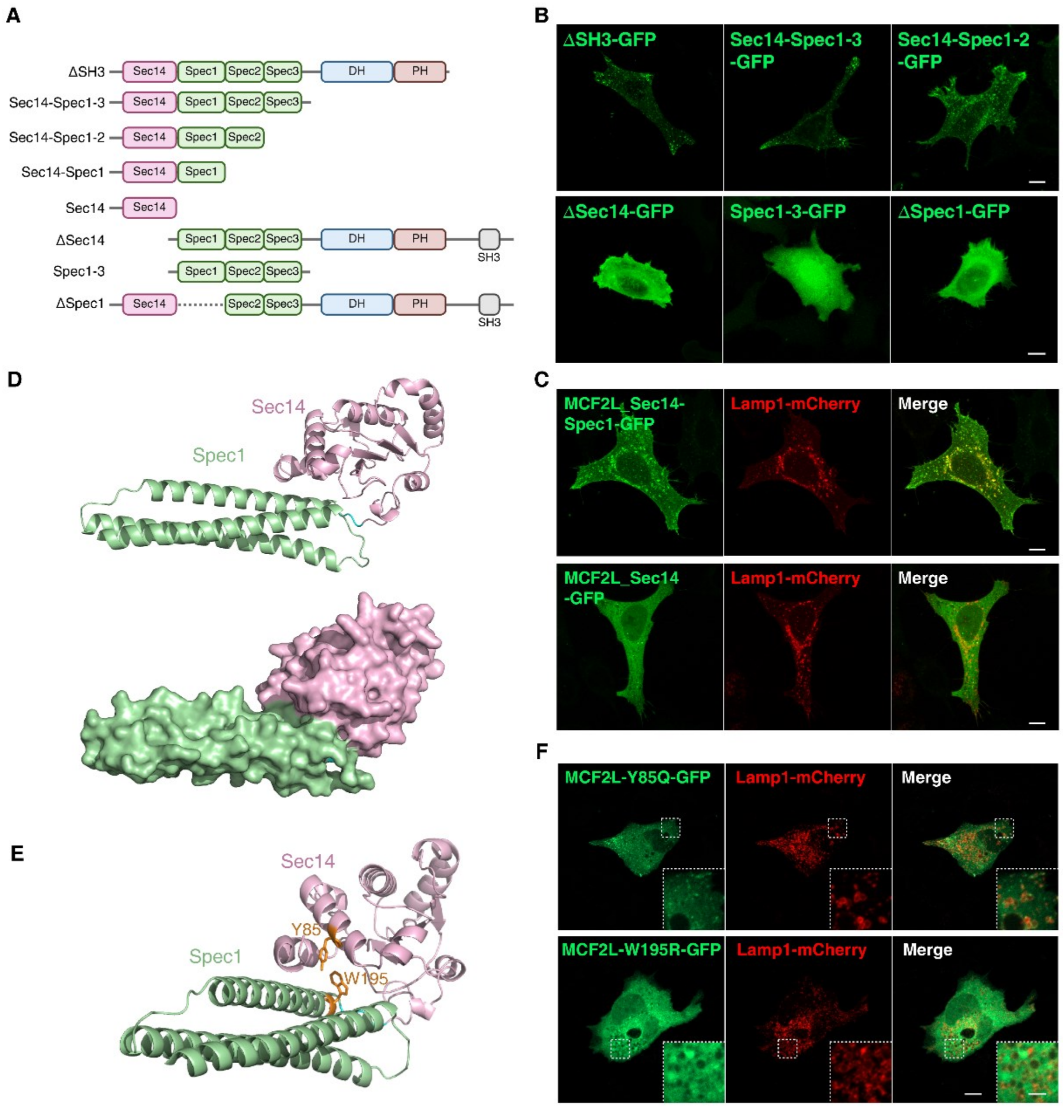


#### Fig S4. Mapping of key regions in MCF2L required for its association with lysosomes

**A.** Schematic diagram showing the MCF2L truncation or deletion mutants generated in this study.

**B.** Localization patterns of the MCF2L truncation/deletion mutants in cells. Loss of either the Spec1 or Sec14 domain results in diffused cytosolic distribution. Scale bar: 10 μm.

**C.** The Sec14-Spec1 tandem associates with Lamp1-mCherry-labeled vesicles, whereas the Sec14 domain alone remains diffused. Scale bar, 10 μm.

**D.** AlphaFold3-predicted structure of the MCF2L Sec14-Spec1 tandem, presented as the ribbon diagram (upper panel) or the surface model (lower panel), showing that Sec14 and Spec1 form a structurally coupled supramodule. Sec14 domain (pink), Spec1 domain (light green).

**E.** Ribbon diagram showing that Y85 and W195 are located in the hydrophobic coupling interface of the Sec14-Spec1 supramodule.

**F.** Confocal images showing that the MCF2L-Y85Q and MCF2L-W195R mutants lose their lysosome association ability. Scale bar, 10 μm; magnified inset, 1 μm.

### Figure S5

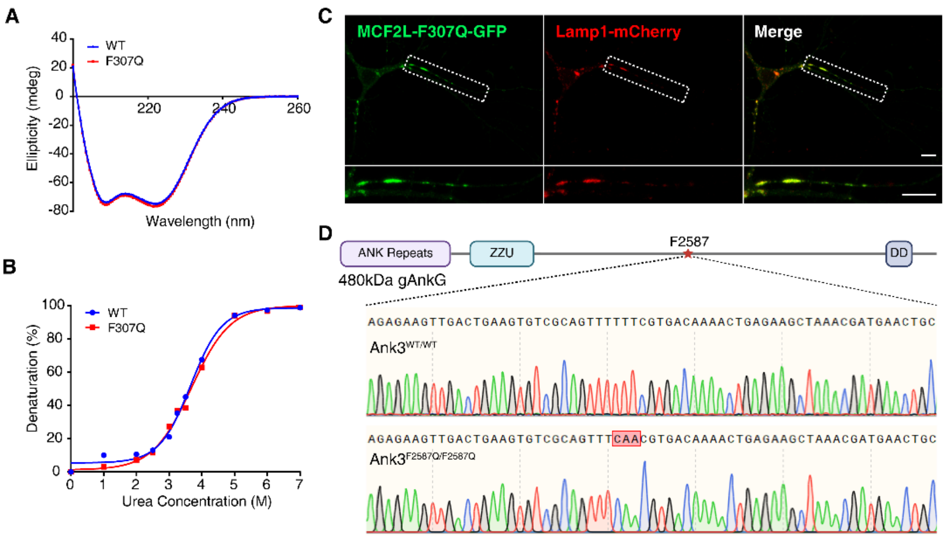


#### Fig S5. Validation of MCF2L mutant protein.

**A.** Circular dichroism spectra of WT and F307Q mutant of MCF2L_Spec1, showing that the mutation does not alter the folding of the protein.

**B.** Urea-dependent denaturation curves of WT and F307Q MCF2L_Spec1, showing that the mutation does not alter the stability of the protein. The data points were derived by recording the ellipticity value of each CD spectrum at 222 nm for each sample at the specified urea concentration.

**C.** MCF2L F307Q mutant does not affect its association with lysosome. Same cell as shown in Fig. 2E, lower panel. Scale bar, 10 μm. Dashed boxes are selected for magnification.

**D.** Sanger sequencing confirmation of WT and the homozygous *Ank3*$^{F2587Q/F2587Q}$ mutant mice.

### Figure S6

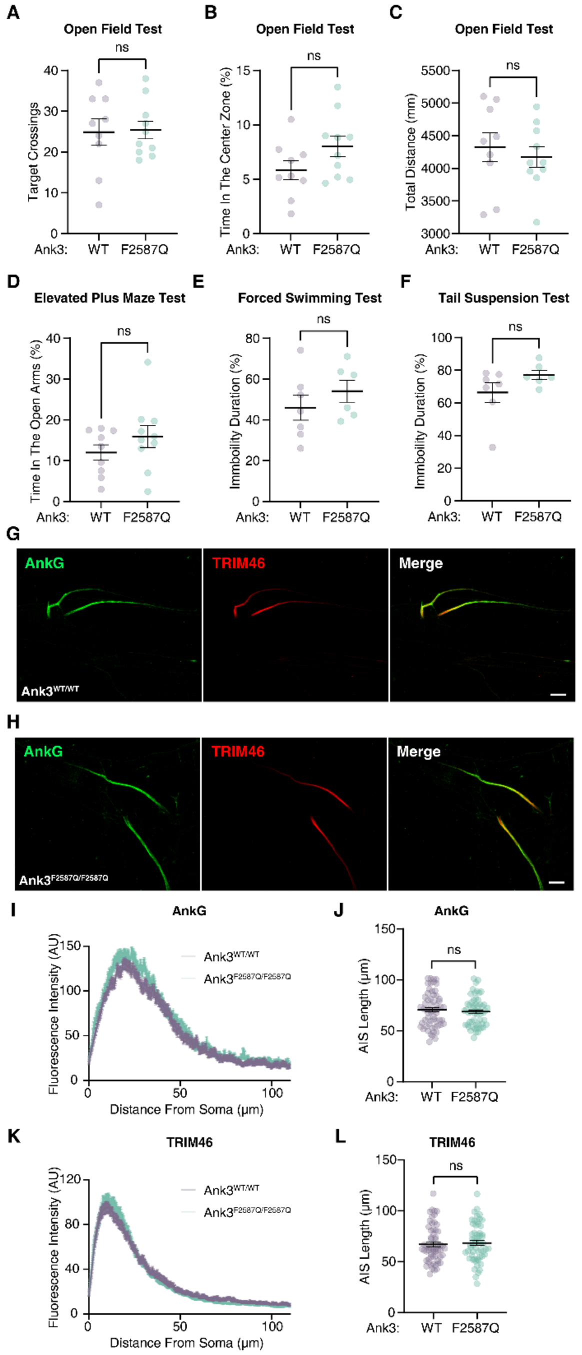


**Fig S6. *Ank3*$^{F2587Q/F2587Q}$ mice behave normally and AIS structural integrity is not affected.**

**A-F.** Behavioral tests revealed no significant differences between adult *Ank3*$^{F2587Q/F2587Q}$ mice and their littermates. Each data point represents an individual mouse. Unpaired two-tailed t test.

**G-H.** Immunofluorescent images of DIV10 hippocampal neurons from *Ank3*$^{WT/WT}$ (**G**) and *Ank3*$^{F2587Q/F2587Q}$ (**H**) mice stained for AnkG and TRIM46. Scale bars, 10 μm.

**I-J**. Quantification of AnkG fluorescence intensity along AIS (**I**) and the AnkG-defined AIS length (**J**) in *Ank3*$^{WT/WT}$ and *Ank3*$^{F2587Q/F2587Q}$ neurons. Mean ± SEM, three independent experiments; n=60 and 57 neurons, respectively. p=0.4945; ns, not significant; two-tailed unpaired Student's t test.

**K-L.** Quantification of TRIM46 fluorescence intensity along AIS (**K**) and the TRIM46-defined AIS length (**L**). Mean ± SEM, three independent experiments; n=60 and 57 neurons, respectively. p=0.5618; ns, not significant; Mann-Whitney test.

### Figure S7

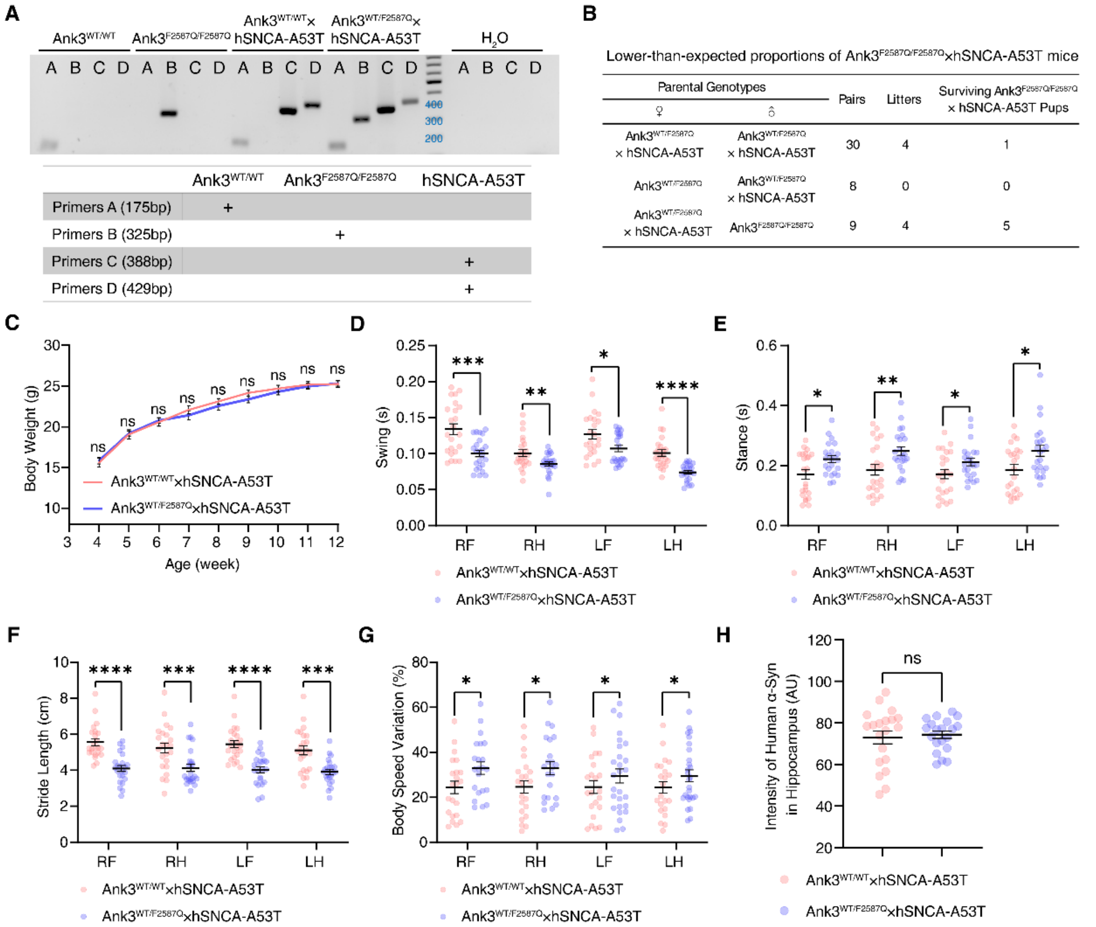


**Fig S7. Additional validation of mice strains and phenotypes for studies related to Fig. 3.**

**A.** Genotyping of the indicated mice by PCR, using pure water sample as the negative PCR control. Primer pairs and expected product sizes are listed.

**B.** Low-yield (non-Mendelian inheritance) in breeding of *Ank3*$^{F2587Q/F2587Q}$×hSNCA-A53T mice. Genotypes of parental pairs, numbers of litters obtained, and surviving pups are listed.

**C.** Body weight curves of male *Ank3*$^{WT/WT}$×hSNCA-A53T and *Ank3*$^{WT/F2587Q}$×hSNCA-A53T mice showing no significant difference between the two mice groups; n>10 mice for each group. ns, not significant, Mann-Whitney test.

**D-G.** CatWalk gait analysis parameters showing differences in swing (**D**), stance (**E**), stride length (**F**) and body speed variation (**G**) between *Ank3*$^{WT/WT}$×hSNCA-A53T and *Ank3*$^{WT/F2587Q}$×hSNCA-A53T mice. Mean ± SEM, three independent experiments; n=23 trials. *p<0.05, **p<0.01, ***p<0.001, ****p<0.0001, Mann-Whitney test.

**H.** Quantification of human α-synuclein intensity in the hippocampus of *Ank3*$^{WT/WT}$×hSNCA-A53T and *Ank3*$^{WT/F2587Q}$×hSNCA-A53T mice. Mean ± SEM, three independent experiments; n=20. ns, not significant, two-tailed unpaired t test.

#### Figure S8

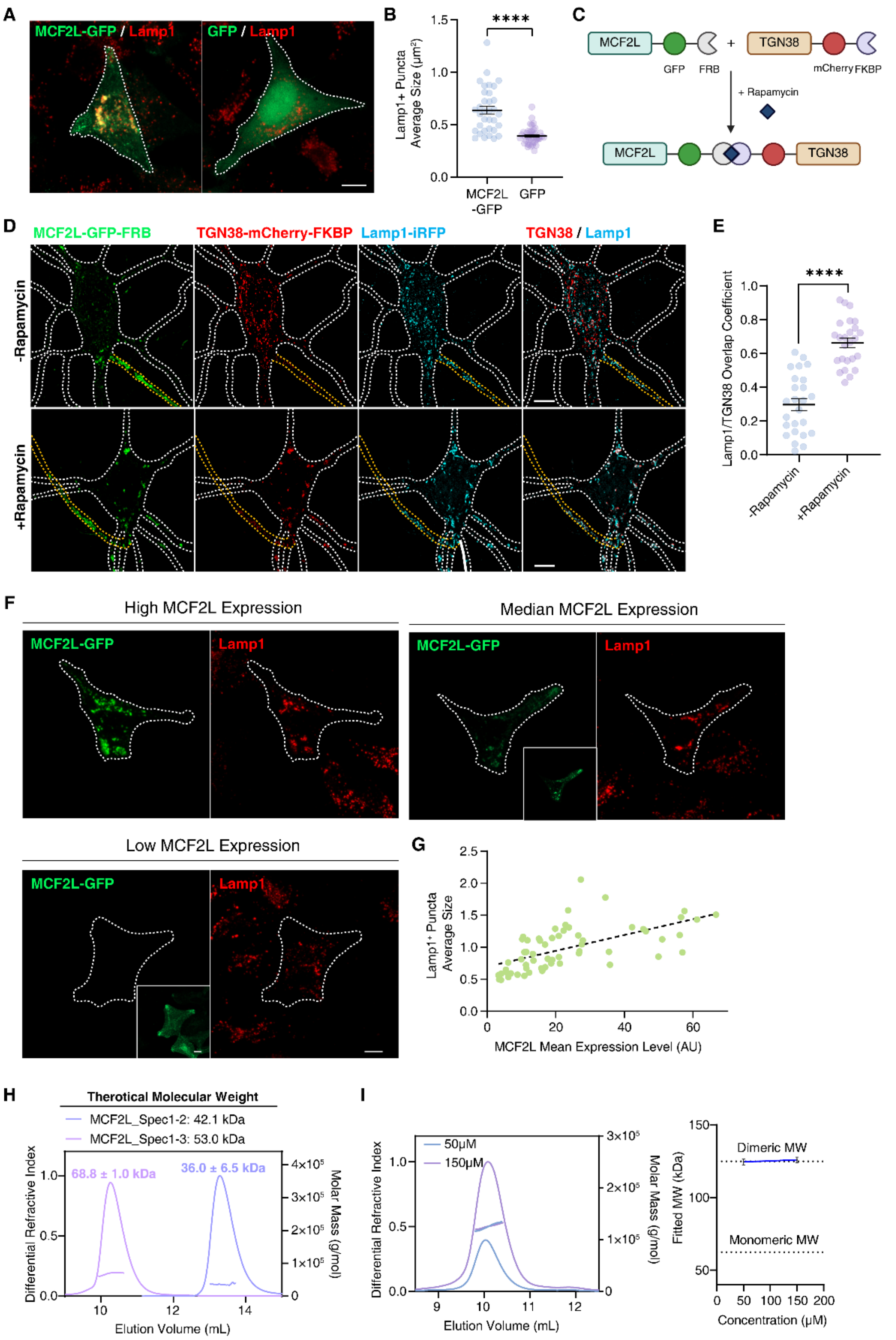


##### Fig S8. MCF2L has organelle-clustering activities.

**A-B.** MCF2L expression markedly increases Lamp1-positive puncta size. **A.** Representative confocal images of HeLa cells expressing MCF2L-GFP or GFP together with Lamp1-mCherry. Scale bar, 10 μm. **B.** Average size of Lamp1-positive puncta in cells expressing MCF2L-GFP or GFP. Mean ± SEM, three independent experiments; n=34 or 41 cells, respectively. ****p<0.0001, two-tailed unpaired t test.

**C**-**E.** Induced coupling of TGN38 to MCF2L increases TGN38 and lysosome colocalization. **C.** Schematic diagram of rapamycin-induced dimerization between MCF2L-GFP-FRB and TGN-mCherry-FKBP. **D.** Hippocampal neurons expressing MCF2L-GFP-FRB and TGN38-mCherry-FKBP with or without rapamycin treatment. White dashed lines indicate the somatodendritic compartment; yellow dashed lines indicate the axon. Scale bar, 5 µm. **E.** Overlap coefficient between TGN38 and Lamp1 in neurons with and without addition of rapamycin. Mean ± SEM, three independent experiments; n=25 neurons. ****$p<0.0001$, two-tailed unpaired t test.

**F**-**G**. MCF2L protein level correlates with Lamp1 puncta size. Confocal images (**F**) of HeLa expressing different amounts of MCF2L and immunostained for Lamp1, with quantifications shown in (**G**). Scale bar, 10 µm. The dashed line indicates a simple linear regression fit.

**H.** SEC-MALS analysis of MCF2L_Spec1-3 and Spec1-2. Theoretical molecular weight (Mw) and determined Mw are shown in the caption or neighboring the curves, respectively.

**I.** *Left*: SEC-MALS analysis of MCF2L_Spec1-3-R518A/S521E-GCN4di at 50 µM and 150 µM. *Right*: Experiment derived Mw at the two concentrations match the expected Mw of a dimer, mean ± SEM.

#### Figure S9

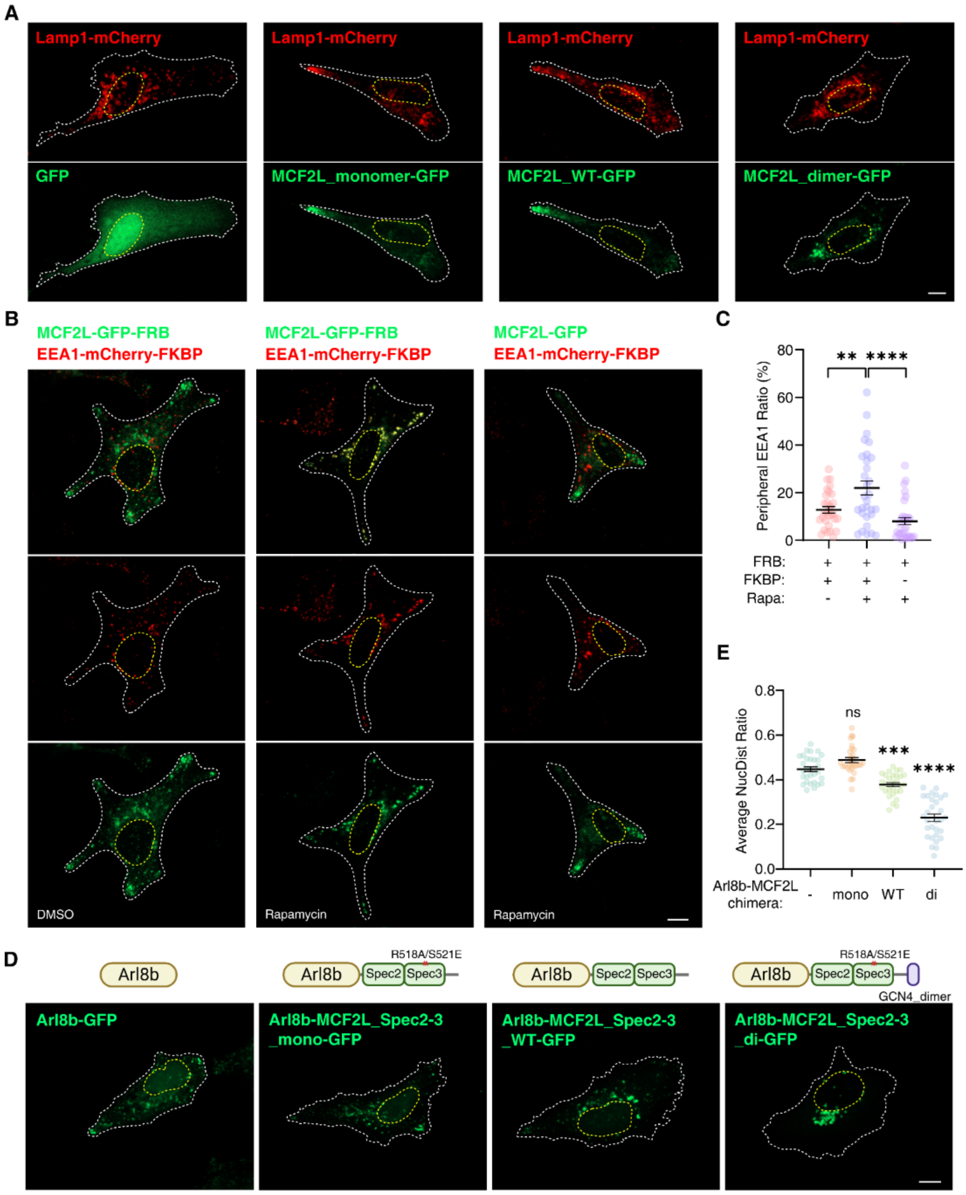


**Fig S9. Lysosome clustering regulates cellular lysosomal positioning.**

**A**. Images of individual fluorescent channels corresponding to Fig. 4G.

**B-C.** Artificial tethering of EEA1-positive endosomes to MCF2L via the rapamycin-inducible FKBP-FRB system significantly increases the proportion of EEA1-positive vesicles at the cell periphery. **B**, representative images, Scale bar: 10 μm. **C**, quantification of EEA1 peripheral distribution. Mean ± SEM, three independent experiments; n=30 cells. **p=0.0065, ****p<0.0001, Mann-Whitney test.

**D-E.** Fusion of Arl8b to distinct MCF2L constructs (WT, monomer, and locked-dimer) results in its differential peripheral enrichments. **D**, representative images, Scale bar: 10 μm. **E**, quantification of the peripheral distribution for each construct. One-way ANOVA followed by Tukey's post hoc test was used; n=30 cells. ***p=0.0005, ****p<0.0001.

### Figure S10

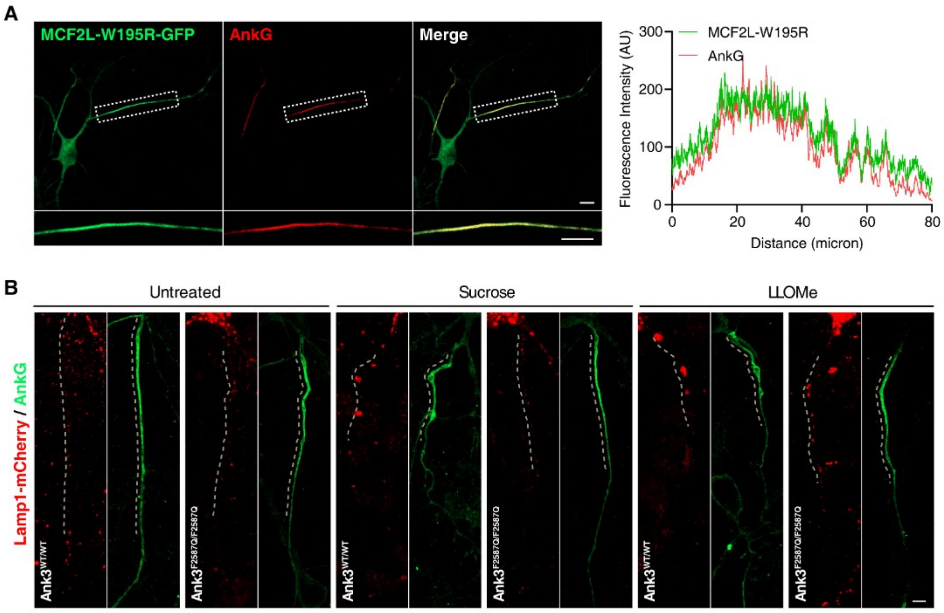


**Fig S10. MCF2L-W195R mutant still localizes to AIS.**

**A.** Confocal images showing that MCF2L-W195R mutant, which is defective in lysosome association, is still enriched at the AIS and well-colocalized with AnkG in hippocampal neurons. Scale bars: 10 μm. Dashed boxes are selected for magnification and line-scan analysis.

**B.** Images of individual fluorescent channels corresponding to Fig. 8E.

#### Movie Legends

##### Movie 1

Confocal live imaging of a DIV8 mouse hippocampal neuron distal axon co-transfected with MCF2L-GFP and Lamp1-mCherry, showing co-transport of the two proteins along the axon.

##### Movie 2

HeLa cells co-expressing Lamp1-mCherry-FKBP and Lamp1-GFP-FRB. Addition of 1 μM rapamycin (t = 0) triggers rapid lysosomal clustering and redistribution. Related to Fig. 5C.

##### Movie 3

SIM live imaging of a DIV8 mouse hippocampal neuron distal axon co-transfected with MCF2L-GFP and Lamp1-mCherry, showing two distinct but interconvertible Ly/LE populations undergoing continuous exchange. Related to Fig. 6E.

##### Movie 4

Same as Movie 3 but showing a different example. Related to Fig. 6E.

**Table S1: Statistics of Data Collection and Model Refinement.**

| | MCF2L/AnkG |
|---|---|
| **Data collection** | |
| X-ray source | SSRF |
| Space group | $P3_121$ |
| Unit cell parameters (Å) | ***a***= ***b***=132.9; ***c***=151.8 |
| Resolution range (Å) | 50-2.90 (2.95-2.90) |
| No. of unique reflections | 34,946 (1,745) |
| Redundancy | 19.8 (20.6) |
| **I/*σ*** | 18.2 (1.2) |
| Completeness (%) | 100 (100) |
| $R_{merge}$ (%)[a] | 19.4 (100) |
| $CC_{1/2}$(highest-resolution shell) | 0.78 |
| **Structure refinement** | |
| Resolution (Å) | 50-2.90 (2.98-2.90) |
| $R_{cryst}$[b] / $R_{free}$[c] | 24.0/25.8 |
| r.m.s.d bonds (Å) / angles (°) | 0.012/1.22 |
| Average B-factors[d] | 72 |
| No. of atoms | |
| Protein atoms | 2825 |
| Water molecules | 11 |
| Ramachandran plot[d] | |
| favored regions(%) | 92.4 |
| allowed regions(%) | 7.6 |
| outliers(%) | 0 |

Numbers in parentheses represent the value for the highest resolution shell.

a. $R_{merge} = \Sigma\ |I_i - \langle I \rangle| / \Sigma I_i$, where $I_i$ is the intensity of measured reflection and $\langle I \rangle$ is the mean intensity of all symmetry-related reflections.

b. $R_{cryst}=\Sigma||F_{calc}| - |F_{obs}||/\Sigma F_{obs}$, where $F_{obs}$ and $F_{calc}$ are observed and calculated structure factors.

c. $R_{free}= \Sigma_T||F_{calc}| - |F_{obs}||/\Sigma F_{obs}$, where T is a test data set of about 5% of the total unique reflections randomly chosen and set aside prior to refinement.

d. B factors and Ramachandran plot statistics are calculated using MolProbity.
